# Single nuclear transcriptional signatures of dysfunctional brain vascular homeostasis in Alzheimer’s disease

**DOI:** 10.1101/2021.10.27.465860

**Authors:** Stergios Tsartsalis, Nurun Fancy, Amy M. Smith, Combiz Khozoie, Xin Yang, Karen Davey, Nanet Willumsen, Aisling McGarry, Robert C. J. Muirhead, Stephanie Debette, David R. Owen, Paul M. Matthews

## Abstract

Brain perfusion and normal blood brain barrier integrity are reduced early in Alzheimer’s disease (AD). We performed single nucleus RNA sequencing of vascular cells isolated from AD and control brains to characterise pathological transcriptional signatures. We found that endothelial cells (EC) are enriched for expression of genes associated with susceptibility to AD. EC transcriptional signatures identified mechanisms for impaired β-amyloid clearance. Evidence for immune activation was found with upregulation of interferon signalling genes in EC and in pericytes (PC). Transcriptional signatures suggested dysregulation of vascular homeostasis and angiogenesis with upregulation of pro-angiogenic signals (*HIF1A*) and metabolism in EC, but downregulation of homeostatic growth factor pathways (VEGF, EGF, insulin) in EC and PC and of extracellular matrix genes in fibroblasts (FB). Our genomic dissection of vascular cell risk gene enrichment suggests a potentially causal role for EC and defines transcriptional signatures associated with microvascular dysfunction in AD.

## Introduction

Alzheimer’s disease (AD) is the most common form of dementia^1^, characterized by extracellular deposits of toxic forms of β-amyloid (Aβ) protein, intracellular neurofibrillary tangles (NFTs) and neurodegeneration. Large-scale genomic association studies have suggested specific molecular processes responsible for susceptibility to disease^2–4^. The non-neuronal cells in which these genes are predominantly expressed are candidates for early “causal” roles in the initiation of the pathological cascades of AD^5^.

Brain microvasculature appears to play a major role in AD pathophysiology^6–8^. Cells constituting the blood brain barrier (BBB) contribute to the clearance of Aβ and other toxic species from the central nervous system (CNS) and allow the selective exclusion of potentially inflammatory or toxic blood proteins from the brain and control of immune cell trafficking^9^. Vascular pericytes are responsible for regulating brain perfusion and contribute to the regulation of endothelial permeability and immune activation^7,10^. Multiple *in vivo* imaging and *post mortem* neuropathological studies, as well as studies of preclinical models, provide evidence for impaired regulation of cerebral blood flow and maintenance of the integrity of the blood brain barrier (BBB) in early AD^11–14^. Recent work has begun to elucidate the transcriptional mechanisms^15–17^.

We have performed an integrated analysis of our own single-nuclei RNA sequencing (snRNAseq) data with that from a previously published dataset^18^ to quantitatively define the enrichment of brain microvascular cells for the expression of AD risk genes as a test of their potential causal contributions to disease genesis^5^. We then explored the functional roles of AD risk genes by assessing functional enrichment of genes co-expressed with them in vascular cells. Differential expression and gene co-expression analyses allowed characterisation of specific genes and pathways altered in AD. A cell-cell communication analysis further defined signalling mechanisms supporting vascular homeostasis and angiogenesis that are impaired with AD. Together, our results provide a transcriptomic mechanistic description for major features of the vascular pathophysiology observed *in vivo* with AD.

## Results

### Endothelial cells are enriched in genes associated with genetic risk for AD

Our analyses were based on data from 57 different brain samples from donors with AD (n=31) or non-diseased controls (NDC, n=26). Fluorescence-activated sorting (FACS) of nuclei isolated before snRNAseq removed neuronal and oligodendrocyte nuclei to achieve a better representation of the less abundant brain cell types of interest. Data was integrated using LIGER^19^ and clustered with UMAP^20^ (Figure 1A). AD and NDC donor nuclei and nuclei from different datasets, brain regions and sexes were well-mixed after integration (Figure S1). Nuclei numbers did not differ significantly between AD and the NDC samples. Feature plots of canonical cell markers identified major brain cell types in the integrated dataset (Figure S2). Endothelial cells (EC) expressed marker genes *FLT1, VWF, NOSTRIN (*Figure S3A)*, CLDN5* and *IFI27*^17,21^ (Figure 1C). Specific expression of *COL1A1, COL12A1, COL6A1* (Figure S3B) and *COL5A1* was used to identify fibroblasts (FB) (Figure 1C). A separate, heterogeneous cluster of vascular mural cell nuclei expressed *PDGFRB, RGS5 and GRM8 (*characteristic of PC^17^) (Figure S3C) and *ACTA2* and *MYH11* (highly expressed in smooth muscle cells (SMC)^17^) (Figure 1C). To distinguish PC from SMC nuclei, we re-clustered the EC, FB and vascular mural cell (PC and SMC) nuclei from the total dataset (Figure 1B) to separate those expressing high levels of *ACTA2* and *MYH11* with very low levels of *RGS5* and *GRM8* (corresponding to SMC) from those expressing high levels of *RGS5* and *GRM8* with very low levels of *ACTA2* and *MYH11* (corresponding to PC) (Figure 1C and S4). We confirmed our cluster annotations by demonstrating significant mutual overrepresentations of our cluster markers and those reported previously in human^17^ (Figure S5A) and mouse^17,21^ (Figure S5B) single nuclei or single cell RNA sequencing studies. To further characterize the identity of the FB population, we tested the overrepresentation of previously described meningeal and perivascular FB markers^15^ and found that our FB markers were more significantly enriched in perivascular fibroblast markers (Fisher’s exact test (FET) for overrepresentation: perivascular FB markers, p = 2.82×10^-77^; meningeal FB markers, p = 6.36×10^-19^).

**Figure 1.**
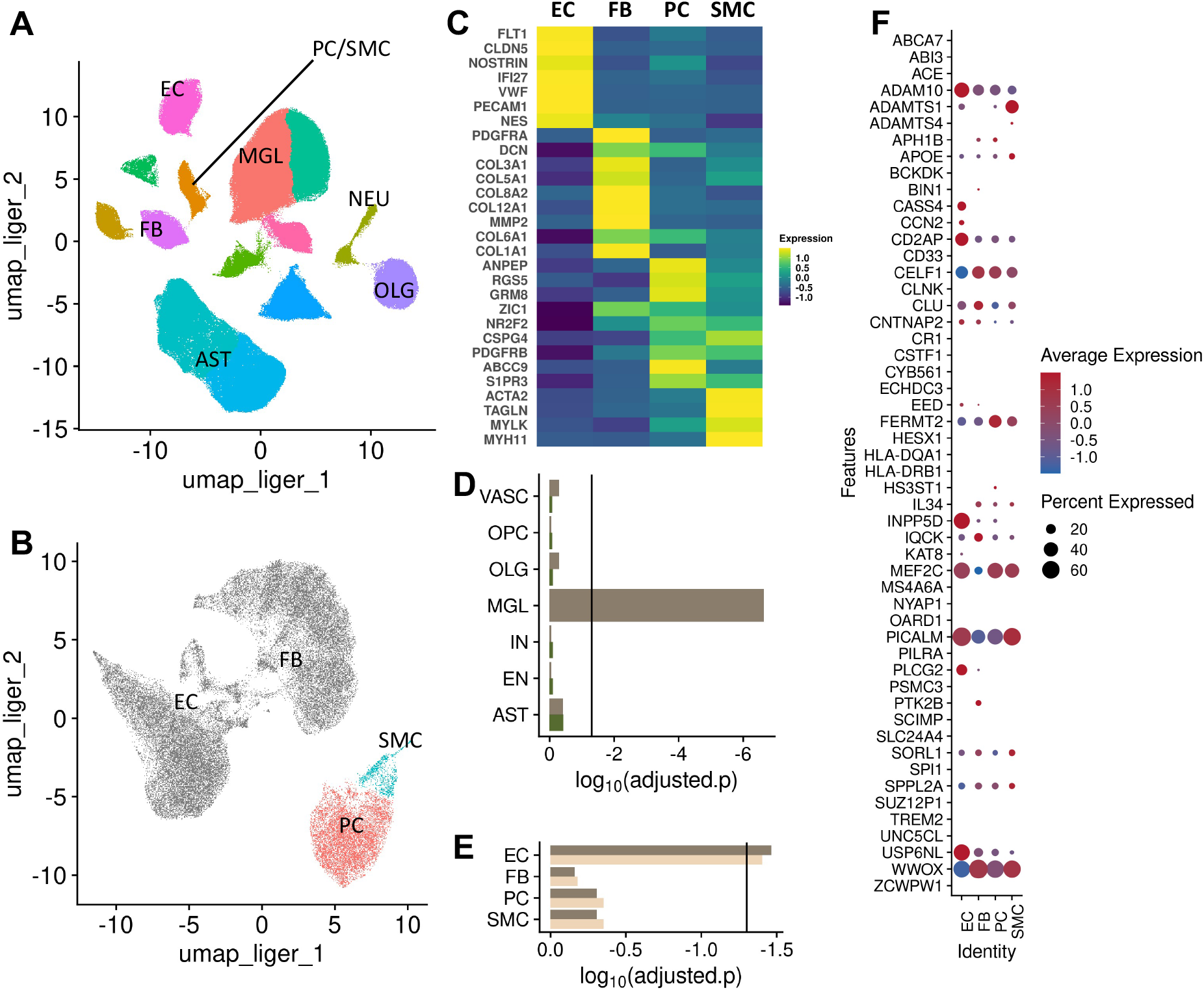
Characterisation of cell-type specific transcriptomes and their relative enrichment in Alzheimer’s disease risk genes. (A) UMAP plot of the integrated snRNAseq dataset from 57 brain samples. (B) UMAP plot after re-integration and clustering of the EC, FB, PC and SMC nuclei in (A), allowing discrimination between PC (red) and SMC (cyan) nuclei (EC and FB, grey). (C) Heatmap of the average scaled expression of representative marker genes for each cluster. (D) MAGMA.Celltyping enrichment of brain nuclei in genomic loci associated with genetic risk for AD. The bars correspond to the log_10_ p value of the enrichment (dark brown, line indicates significance threshold adjusted for all cell types). Enrichment of vascular nuclei is reduced after controlling for genes enriched in microglia (dark green) (F) MAGMA.Celltyping AD risk gene enrichment of nuclei of the brain vasculature (dark brown bar, line indicates significance threshold adjusted for vascular cell types). Enrichment is not changed substantially after controlling for the enrichment of genetic loci associated with white matter hyperintensities (WMH) (light brown). (H) Dot plot of the average scaled per cluster expression of genes previously associated with genetic risk for AD (size, percentage of nuclei per cluster with >1 count; colour scale, average scaled gene expression). Abbreviations: AST, astrocytes; EC, endothelial cells; FB, fibroblasts; MGL, microglia; NEU, neurons; EN, excitatory neurons; IN, inhibitory neurons; OLG, oligodendrocytes; PC/SMC, pericytes and smooth muscle cells.

Well-annotated genes associated with genetic risk of AD^2–4^ were expressed in nuclei from all four vascular cell types (Figure 1F): 52/61 AD risk genes tested were found in at least one of the vascular cells studied, although less than half of these genes were expressed in 5% or more of nuclei (EC, 21/61; FB, 21/61; SMC, 17/61; PC, 19/61). 14/61 of these genes were expressed in at least 5% of the nuclei across all *four* cell types (*ADAM10, APOE, CD2AP, CELF1, CLU, CNTNAP2, FERMT2, IQCK, MEF2C, PICALM, SORL1, SPPL2A, USP6NL, WWOX*).

We employed MAGMA.Celltyping to test for the significance of the enrichment of vascular nuclei across the larger set of genomic loci associated with AD^5^. First, we analysed a dataset that included all the canonical cell types of the brain (Figure S6-S8). This showed that the AD risk gene expression enrichment is greatest in microglia, as has been reported previously^5^ (Figure 1D). Vascular cells also were relatively enriched, albeit less than microglia. To partition enrichment amongst the individual vascular cell types, the analysis was repeated with vascular cell data alone. Only EC were significantly enriched for expression of AD risk genes (Figure 1E). Brain small vessel disease and MRI brain white matter hyperintensities (WMH) are associated with risk of AD^22,23^. To test whether risks for small vessel disease were responsible for the EC enrichment for AD genetic risk, we re-assessed enrichment for the latter after statistically controlling for WMH risk gene expression^24^. The results remained virtually unchanged (Figure 1E): the genetic risk for AD associated to the EC transcriptome thus appears to be independent of that for WMH. However, when the analysis was repeated after controlling for the microglial enrichment, the vascular enrichment largely disappeared, suggesting that similar AD risk gene sets are enriched in microglia and vascular (Figure 1D).

### Differential gene expression (DGE) identifies transcriptional signatures of dysfunctional angiogenesis in AD

We employed a mixed-effects model in MAST^25^ to discover genes differentially expressed in AD relative to NDC for each of the cell types. Greater numbers of genes were downregulated (90 genes, EC; 47 genes, FB; 47 genes, PC; FDR 0.1), than upregulated (73 genes, EC; 25 genes, FB; 25 genes, PC) in EC, FB and PC (Figure 2A-C). We did not find significant differentially expressed genes (DEG) in the SMC.

**Figure 2.**
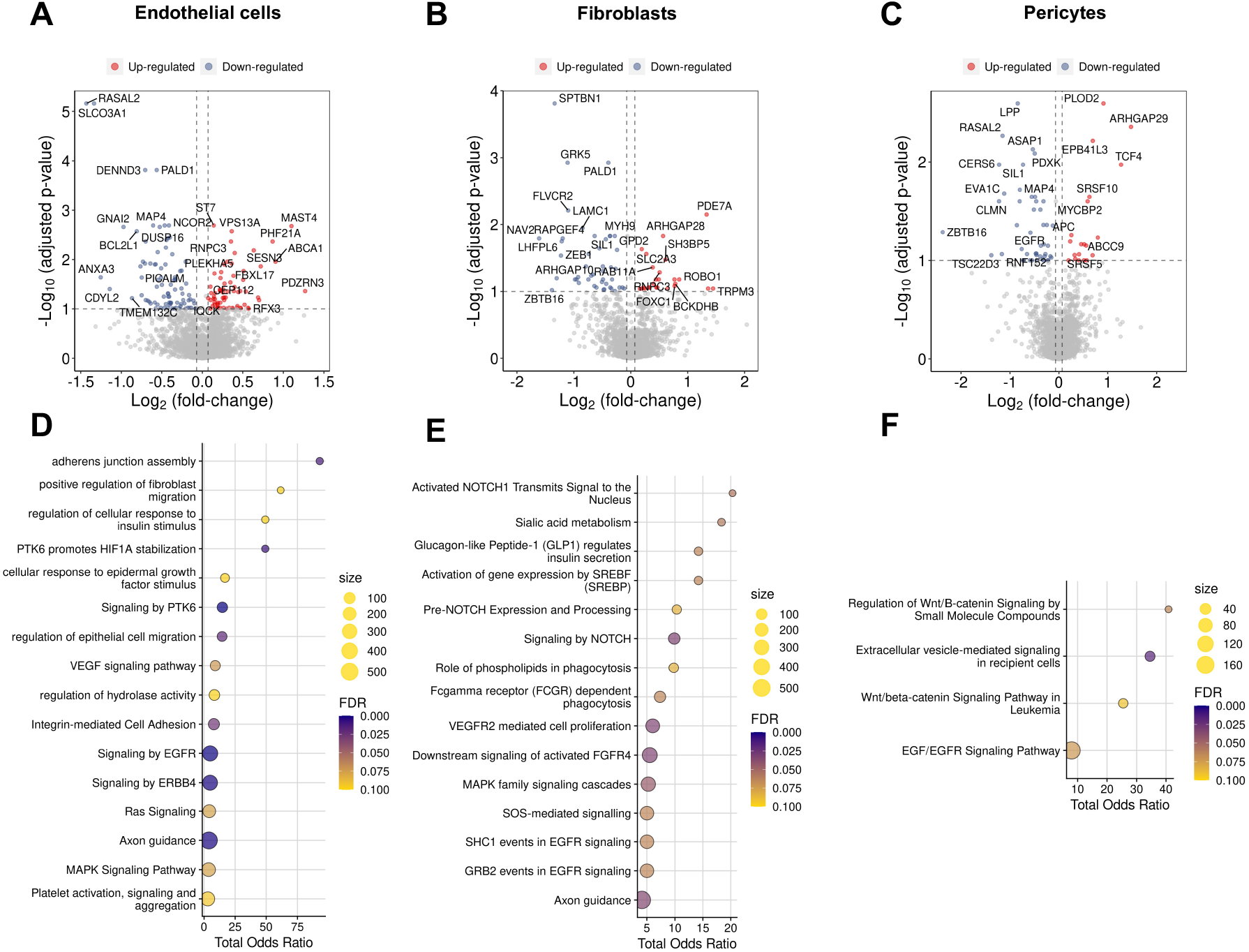
Alzheimer’s disease is associated with dysregulation of vascular homeostasis. Volcano plots showing genes differentially expressed in AD relative to NDC in (A) EC, (B) FB and (C) PC. Corresponding dot plots of the functional enrichment analysis on the DEG (D-F, (dot size, functional enrichment gene set size; colour, FDR).

### Pathological angiogenic transcriptional signatures in AD

Proangiogenic *HIF1A* was overexpressed in EC in AD (Figure 2A). However, the expression of multiple functionally related genes (e.g., *SPRED2, SHC2, KSR1, RASGRF2, DAB2IP, RASAL2, DUSP16, VCL* and *EGFR)* involved in VEGFR2, EGFR and insulin receptor-mediated pathways were downregulated. Pathological angiogenic gene expression signatures also were found in FB with downregulation of the expression of VEGF, FGF, EGF and IGF pathway genes (including *SPRED2*, *DAB2IP* and *SPTBN1*), *DTX2*, a regulator of Notch signalling^26^ and the sialyltransferase gene, *ST3GAL6* (Figure 2B, E). Gene expression in PC highlighted a strikingly mixed angiogenic signature with upregulation of the angiogenic Wnt/β-catenin signalling pathway and downregulation of EGF/EGFR signalling pathway genes including *RPS6KA2, ASAP1, MEF2D* and *EGFR* (Figure 2C, F).

Clues to additional mechanisms responsible for loss of BBB integrity in AD were found with the differentially expressed gene signatures. For example, genes contributing to adherens junction assembly (*VCL, TBCD* and *PIP5K1C*) were variably differentially expressed in EC (*VCL* and *TBCD* were downregulated, whereas *PIP5K1C* was upregulated). As noted above, Wnt/β-catenin pathway genes, including *TCF4* and *APC,* expression of which also support blood brain barrier integrity^27^, are significantly upregulated with AD in PC. Finally, *LAMC1,* encoding for the gamma laminin subunit, was significantly downregulated in FB.

### Differential expression of risk genes with functional roles in amyloid processing and immune response in AD

Risk genes with functional roles related to amyloid processing were differentially expressed in AD. *ADAM10*, which encodes the constitutive α-secretase that governs non-amyloidogenic pathway β-amyloid precursor protein processing, was significantly upregulated in PC. *PICALM*, encoding a clathrin assembly protein modulating clearance of Aβ, was downregulated in EC. Risk genes related to immune responses also were differentially expressed with AD. Increased expression of the inhibitory complement receptor *CD46* gene and decreased expression of *IRAK3,* which encodes an IL-1 receptor associated kinase were found in PC. In EC, we also found increased expression of *IFITM3,* which regulates interferon pathway inflammatory responses^28^ and can potentiate gamma secretase activity^29^.

### Co-expression modules for angiogenesis, lysosomal processing and interferon activation are differentially regulated with AD

To identify gene co-expression modules differentially expressed with AD, we first performed gene co-expression network analyses separately for EC, FB and PC pooled across AD and NDC (MEGENA^30^). We then determined which gene co-expression modules were differentially associated with AD (limma^31^). SMC were not included in this analysis because of the relatively low number of nuclei available for analysis and consequent sparse co-expression representation. Our results defined cell-specific gene regulation pathways associated with AD.

### Reduced expression of co-expression modules enriched for angiogenesis and vascular homeostasis

Although fold-changes varied, expression of modules enriched for multiple angiogenic or vascular homeostasis pathways were generally decreased with AD, most prominently for EC (Figure 3), e.g., Module 119, which is enriched in the EGF/EGFR signalling pathway (e.g., including *EGFR*, *IQSEC1*, *INPP5D*, *NEDD4* and *IQGAP1*) and several hub genes (e.g., the G-protein activator and adhesion G protein coupled receptor genes, *DOCK9* and *ADGRL4,* respectively, and the hypoxia inducible transcription factor, *EPAS1)* with roles in angiogenesis^32,33^. Expression also was reduced for module 47, which is enriched in insulin-like growth factor 1 receptor (IGF1R) (*IGF1R*, *PSMD5*, *PSMD1*, *TSC1*, *RASAL2*) and Ras signalling cascade (*PSMD5*, *PSMD1*, *RASAL2*) genes. Expression of several genes in module 47 with recognised functional roles in angiogenesis (e.g., *RASAL2* and *PALD1*^34^) were significantly independently downregulated in AD.

**Figure 3.**
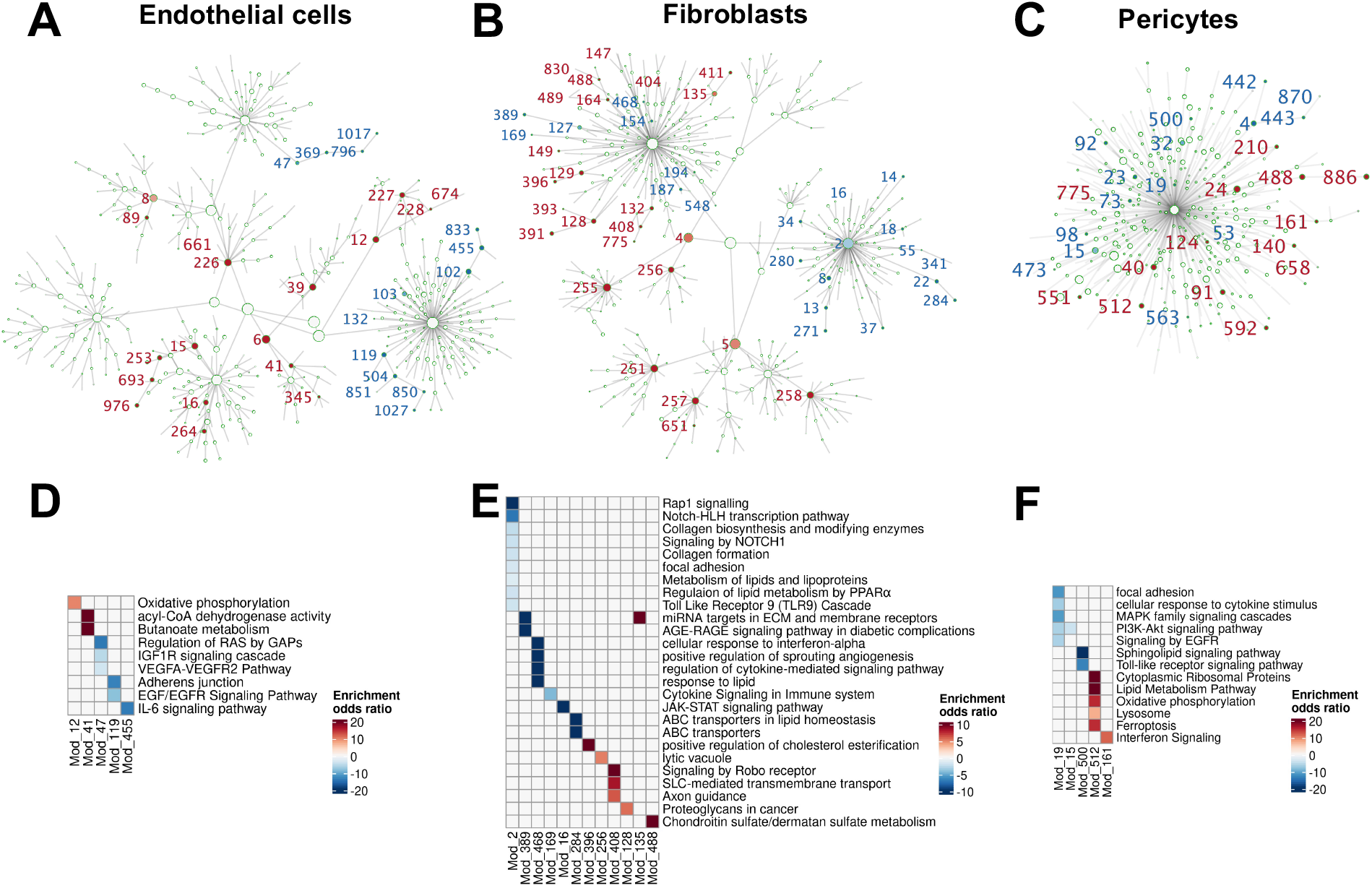
Gene co-expression module hierarchy for (A) EC, (B) FB and (C) PC. Modules that are significantly differentially enriched in AD relative to NDC nuclei are represented as points in the graph (red, upregulated modules with AD; blue, downregulated modules). (D-F) Heatmaps of odds ratios from the functional enrichment analyses for significantly differentially enriched modules in EC (D), FB (E) and PC (F) (red, upregulated; blue, downregulated).

While the fold-changes were relatively low, FB modules functionally related to vascular homeostasis also were significantly downregulated, e.g., Module 2, which was significantly enriched in NOTCH signalling genes (Figure 3E), including *NOTCH1* and *NOTCH2*, as well as the regulators of NOTCH expression^26^, *ARRB1* and *DTX2,* both of which also were independently significantly differentially reduced in expression with AD. Module 2 also is significantly enriched in genes individually downregulated in AD (FET adjusted p=1.28×10^-8^). Some of them (*SPRED2*, *DAB2IP*, *ARRB1*, *SPTBN1)* encode for proteins that are downstream components of several growth factor signalling pathways (e.g., FGFR1-4, VEGFR2, EGFR) or genes for components (*COL5A1* and *COL1A2*) of the extracellular matrix (ECM)^35^. Angiogenesis pathway enriched modules were downregulated in PC, as well, e.g., module 19 (Figure 3F), in which several genes showing individually significantly reduced expression in AD (*EGFR*, *ZBTB16*, *IRAK3*, *TMTC1*, *MAOA*) and genes involved in EGFR signalling (*EGFR, MAPK1, FOXO3*) are found or module 15, which is enriched in PI3K-Akt signalling pathway genes (*ANGPT2, COL6A2*, *DDIT4*, *COL6A1*, *BCL2*, *PDGFA*, *PPP2R3B*, PPP2R3A) genes^36^. A recent report has shown the *Angpt2* knock out potentiates BBB leakage in a preclinical Aβ mouse model^37^.

However, despite the decreased expression of modules enriched for many angiogenic pathways central to angiogenesis with AD, we also found modules enriched in metabolic pathways supporting angiogenesis in EC and PC and extracellular matrix genes in FB, the expression of which was *increased* in AD. Module 661, the top upregulated module in AD, includes *TPI1* that encodes for the triosephosphate isomerase, an enzyme implicated in glycolysis and gluconeogenesis^38^. Module 41, the expression of which is increased with AD, includes genes encoding for the acyl-CoA dehydrogenase that executes the first step of the β-oxidation of fatty acids (*ACAD8, ACADS)* and genes of the butanoate metabolism pathway (*BDH2, ACSM3, and ACADS*)^39^, both of which are highly upregulated in AD. Module 12, also increased in expression with AD, is amongst several in EC that were enriched for pathways coding for proteins of the mitochondrial respiratory chain complexes, including *ATP5PF*, *ATP5PD, ATP6 NDUFB10* and *NDUFS3*. The most highly upregulated module in PC, module 512, was enriched in oxidative phosphorylation genes (*NDUFA4L2, ATP5MC2, ATP5F1D*) and for lipid metabolism pathway genes (*PDHA1*, *PRKAB1*)^40,41^. Fibroblasts play a central role in the development of the basement membrane. Extracellular matrix collagen genes (*COL3A1, COL5A2, COL5A3, COL11A1, COL21A1*), the fibronectin 1 gene (*FN1)*, proteoglycan and glycosaminoglycan metabolism-related genes (*CHSY3, GXYLT2, GPC6, CHST15, TIAM1, KDR, PLCE1, COL21A1, FLNB, ANK3, TP53*) were enriched in modules upregulated with AD in FB, e.g., in branches of modules 128, 135, 164, 408, 488 and 775. Modules 408 and 391, also upregulated with AD in FB, were enriched in solute carrier genes involved in nutrient and metabolite transfer across the blood brain barrier^42^.

### Increased expression of APOE and lysosomal pathway enriched modules in FB and PC with AD

Module 396, the most highly upregulated module in FB with AD, was enriched in the AD risk gene *APOE* and other cholesterol metabolism-related genes (*e.g., AGT),* as well as genes with individually increased expression in AD that are related to pathways for lytic vacuole functions (including *CACNG7*, *VPS28*, *CTSO, RRAGC, NPC2, GAA, HPS4, CTNS, RAB9A, VAMP4, VPS16* and *RAB7A*). Expression of module 512, which is enriched for lipid metabolism (with *PDHA1* and *PRKAB1*) and lysosome (with *PSAP, CLTB* and *CTSF*) pathways, increased in PC, whereas other lipid processing and metabolic gene pathways were downregulated with AD, notably in module 500 (enriched for the sphingolipid signalling pathway genes *AKT3, MAPK14, PLCB1* and *NSMAF*). FB modules 2 (enriched for “regulation of lipid metabolism by PPAR-*α*” and “metabolism of lipids and lipoproteins” pathways) and modules 468 (enriched for “response to lipid”) and 284 (enriched for the “ABC transporters in lipid homeostasis” pathway) also were downregulated with AD.

### Module enrichments suggest pathological amyloid processing and immune responses in AD

There was evidence for upregulation of Aβ production and immune responses with AD. Module 41, which was highly upregulated in EC, contains *PSEN2* and *APH1A*, which encode for components of the gamma secretase^43^. We also found increased expression of interferon-related co-expression modules in EC (module 661, the most highly upregulated modules includes *C2* and *TRIM5* which is a interferon type I-stimulated gene^44^) and in PC (module 161 includes *RNASEL, EIF4A3, AAAS* and *IFIT1*) with AD. The most highly upregulated module in PC (Module 512) is enriched for pathways for ferroptosis (*GPX4, FTH1)*, which promotes release of oxidised lipid species that generate pro-inflammatory damage-associated molecular patterns (DAMPs) able to trigger the innate immune system^45^. However, modules enriched for other immune response pathways were *downregulated* with AD, e.g., EC module 455, which is enriched for IL-6 signalling (*NLK, IL6ST, JAK1*), and PC module 19, enriched for cytokine signalling, and modules 32 and 500, which are enriched for Toll-like response genes. This was most striking for FB, for which the largest number of immune response modules differentially expressed was identified (modules 468 and 154 enriched for interferon-alpha responses, module 2 enriched for the Toll-like receptor 9 (TLR9) cascade, module 169 enriched for cytokine signalling (*IL15, KIT, RAF1)* and module 16 enriched for JAK-STAT signalling), all of which were downregulated with AD.

### Two-layer neighbourhood analysis of risk genes suggests cell-specific mechanisms of AD susceptibility

Cell-specific enrichments for risk genes expression provide insights into the genesis AD^5^. Cell specific co-expression signatures can suggest specific functional roles for the risk genes in susceptibility. To explore those relevant to the cerebral microvasculature, we identified genes having the most direct expression correlations (those within a two-layer neighbourhood, i.e., any gene that is either directly connected to an AD risk gene or through at most one other gene) with AD risk genes in the vascular cell-specific co-expression networks generated from both AD and NDC^30,46^. To discover relationships specifically relevant to disease genesis, we determined the overrepresentation of genes differentially expressed with AD in the neighbourhoods of each GWAS gene in the cell-specific co-expression networks (Figure 4A).

**Figure 4.**
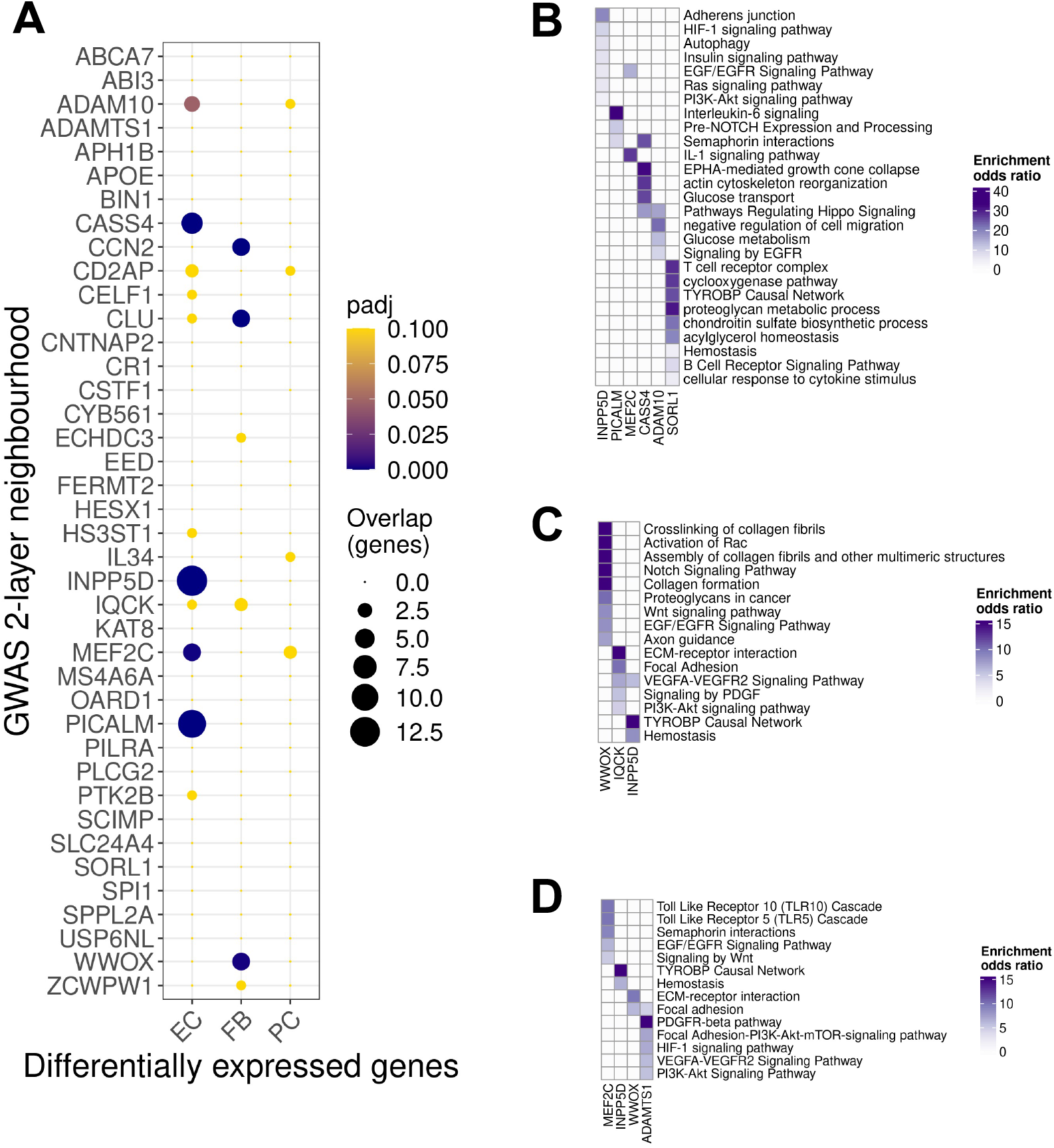
Exploration differentially expressed genes (DEG, AD vs. NDC) in two-layer neighbourhoods of AD risk genes. (A) Dot plot of the overrepresentation of DEG identified in each cluster (abscissa) in AD relative to CTR in the 2-layer neighbourhood of each GWAS gene (ordinate) (dot size, number of the overlapping genes; colour, adjusted p value).B-D) Functional enrichment of prioritized GWAS genes in EC (B), FB (C) and PC (D)(colour scales represent the odds ratio of the enrichment).

AD risk genes *PICALM, SORL1* and *INPP5D* had the largest neighbourhoods (230, 167 and 121 differentially expressed genes, respectively) in the EC co-expression network. The neighbourhood of *PICALM* was enriched in IL-6 signalling genes (*IL6ST, STAT3, JAK1*), as well as semaphorin (*SEMA5A*, *ARHGEF11*, *SEMA6D*, *ITGA1*, *MYH11*, *PLXNC1*) and NOTCH signalling-related genes (*TNRC6C, B4GALT1, TFDP2, POFUT1, MAMLD1, TNRC6A*). The *SORL1* neighbourhood also was enriched in genes for pathways involved in immune response (e.g., T-cell activation, TYROBP causal network and cytokine response-related pathways) and proteoglycan metabolism (chondroitin sulfate biosynthesis and proteoglycan metabolism) pathway genes encoding proteins essential for vascular extracellular matrix formation. The functional enrichment of the *INPP5D* neighbourhood showed an overrepresentation of individual differentially expressed genes that were downregulated in insulin- and EGF/EGFR-signalling pathways (Figure 4B).

While the co-expression neighbourhoods of AD risk genes *WWOX*, *CLU and CCN2* expressed in FB (e.g., including individually differentially regulated genes *ITPR2*, *ROBO1*, *LHFPL6* and *SLC38A1 [WWOX]*; *PALD1, ZBTB46, PDZD2, SIL1* [CLU]; *SPTBN1*, *ZMIZ1*, *KLF7*, *CACNA2D3 [CCN2]*) and PC were enriched for genes involved in a range of functions, the neighbourhood of *WWOX* also was enriched in Notch signalling pathway genes and both this neighbourhood and that of *IQCK* were enriched in ECM-related pathways (Figure 4C). In PC, the neighbourhood of *MEF2C* transcription factor (108 genes) (one of the largest) included genes involved in the EGF/EGFR signalling pathway (*EPS8*, *MEF2A*, *MEF2C*, *PLCE1*, *RAF1*) and Toll-like receptor pathways (e.g. *MEF2A*, *PPP2CB*, *MEF2C* and *RIPK2*) (Figure 4D).

### Pathological growth factor and ECM signalling between vascular cells with AD

We applied CellChat to identify differentially expressed receptor-ligand pairs responsible for cell-cell signalling (both autocrine and paracrine) in AD^47^. 759 potential ligand-receptor pairs implicating 48 distinct signalling pathways were identified amongst genes expressed EC, PC and FB (Figure 5). We first focused on assessment of interaction pairs that involved DEG and pathways enriched in co-expression modules specifically with AD. The analysis provided insight into mechanisms contributing to dysregulated angiogenesis with AD. Several VEGF-related signalling pairs were detected only in NDC. Signalling via INHBA (FB/PC) – ACVR1B and ACVR1A (EC) also was found only in NDC. By contrast, evidence for TGFB3 (EC) – TGFBR1_R2 (FB) and the TGFB3 (EC) – ACVR21B_TGFBR2 (FB) signalling was found only in AD samples.

**Figure 5.**
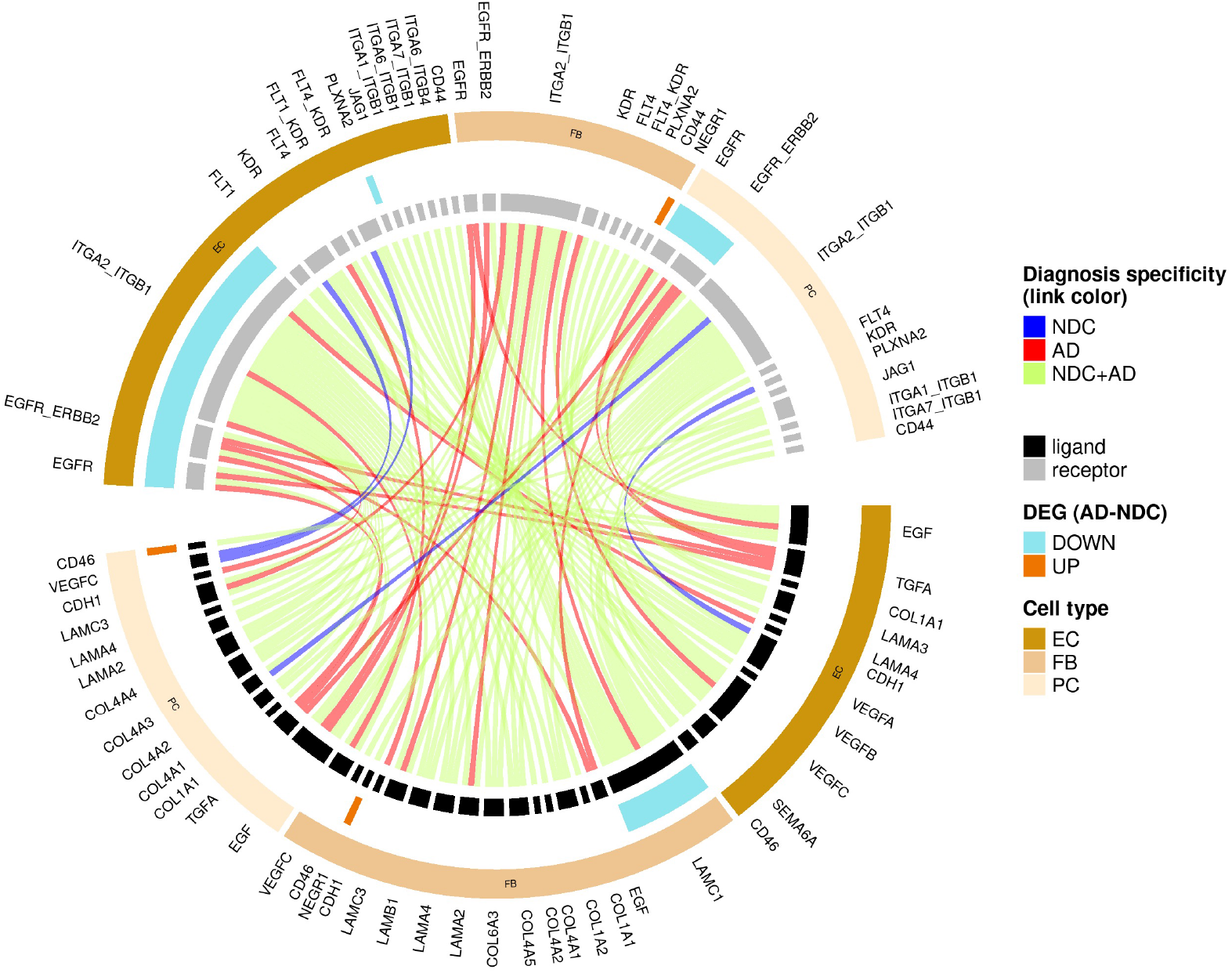
Circular plot of representative results from the cell-cell communication analysis (CellChat v0.5.5). The innermost circle links connect the gene symbols of the ligands with their corresponding receptors. The link colour denotes the diagnosis specificity (i.e. if a ligand-receptor pair was identified only in NDC (blue), only in AD (red) or in both conditions (green). Moving outward, the second track of the plot describes whether the gene corresponds to a ligand (black sectors) or a receptor (grey) in the communication. Genes that were also identified as differentially expressed with AD are highlighted (cyan, downregulated; orange, upregulated). Finally, the cell type where the ligand or the receptor gene is expressed is denoted in the outermost track of the plot.

Growth factor signalling important for angiogenesis and vascular homeostasis also was generally decreased with AD. Receptor expression for the EGF(EC/PC)-EGFR(EC/PC), EGF(EC/PC) - EGFR_ERBB2(PC), NTF3(FB/PC) - NTRK3(FB) communication pairs was reduced. Decreased expression of FB *LAMC1* should reduce autocrine and EC/PC CD44 signalling. Increased expression of *CD46* in PC with AD may modulate NOTCH signalling in an autocrine manner (CD46 (PC)-JAG1(PC))^48^. Autocrine NEGR1(FB)-NEGR1(FB) signalling also increases in AD. Vascular cell interactions with the ECM were reduced with decreased integrin expression on EC affecting multiple ligand-receptor communication pairs: COL1A2 (EC/FB) - ITGB1 (EC), COL1A2 (FB) -ITGA2 (EC), COL4A1 (FB/PC) - ITGA2 (EC) or COL4A2 (EC/FB) - ITGB1 (EC), COL4A4 (PC) - ITGA2_ITGB1(PC) and COL6A3 (FB) - ITGA2_ITGB1 (EC). Both lower EC and PC integrin and FB laminin expression in AD also should reduce the many laminin-ECM interactions identified (LAMC1 (FB) - ITGA1_ITGB1 (EC/PC) - ITGA2_ITGB1 (EC), LAMA3 (EC) - ITGA2_ITGB1(EC), LAMA4 (FB/PC) - ITGA2_ITGB1(EC), LAMB1 (FB) - ITGA2_ITGB1(EC), LAMC1 (FB) - ITGA2_ITGB1 (EC/PC), LAMC3 (EB/PC) - ITGA2_ITGB1 (EC), LAMC3 (FB/PC) - ITGA2_ITGB1 (EC/PC), LAMC1 (FB) - ITGA6_ITGB1(EC/PC), LAMC1 (FC) - ITGA7_ITGB1 (EC/PC)).

### Expression of vascular endothelial angiogenic pathway genes is related to Aβ plaque and phosphorylated Tau (pTau) load

To explore how vascular transcriptomic pathology may evolve over the course of AD, we performed an exploratory analysis of gene expression as a regression of the individual brain regional Aβ and pTau densities in the 24/57 brain samples used in the analyses above for which quantitative IHC was available. We limited this regression analysis to the EC, the most abundant of the cell populations, to minimise Type I errors. We found 28 genes were significantly (adjusted p<0.1) differentially expressed with greater brain regional Aβ plaque density and 75 genes differentially expressed with greater pTau density. Genes showing significant regression with Aβ density in EC were enriched in VEGF signalling pathway components (*CYFIP1*, *PXN*, *CTNNA1*, *CALM1*). By contrast, genes that showed significant regression against pTau were enriched in EGFR, IGF1R and insulin receptor signalling pathways (*PPP2R5E*, *NF1*, *CALM1*, *DUSP16*, *AP2M1*, *RBX1*).

## Discussion

Brain vascular structural pathology and physiological dysfunction is characteristic of both preclinical models expressing brain Aβ and of AD^13,49^. We found that, although EC, SMC and PC all express AD risk genes, EC appear to be uniquely *significantly* enriched amongst vascular cells, suggesting a quantitatively more important role in “causally” mediating AD susceptibility. Our results provided evidence that pathological regulation of angiogenic gene expression contributes to the early vascular impairments in AD: we found upregulation of the pro-angiogenic *HIF1A* and increased expression of mitochondrial oxidative and fatty acid oxidation genes, metabolic drivers of angiogenesis^50^, in conjunction with *downregulation* of vascular developmental and homeostatic pathways involving EGF/EGFR, VEGF/VEGFR and insulin and IGF signalling in EC and PC and reduced expression of laminin and collagen IV genes in FB. The correlation of these expression differences in angiogenic growth factor pathways in EC with both the Aβ plaque and pTau burden suggests that impairments of vascular growth factor pathway signalling worsen progressively through the course of the disease. Analyses of cell-cell communication provided novel evidence for decreased signalling though VEGFR2 and components of the ECM to EC, decreased expression of the LDLR pathway for clearance of Aβ from the CNS by EC, interferon pathway activation and increased inflammatory responses in EC and PC. Together, these processes would be expected to reduce clearance of Aβ while also, via inflammatory activation, increasing pathological Aβ production and processing^29^. Specific evidence for this was found with upregulation in EC of *PSEN2* and *APH1A*, which encode for components of the gamma secretase.

A recent preprint also reported that GWAS risk genes were enriched in EC and vascular mural cells and suggested an “evolutionary shift” of AD risk gene expression from a singular predominance in microglia in the mouse to microglial and vascular cells in humans^15^. Our analysis extends this by showing that, among vascular cells, risk genes associated with AD are enriched significantly in EC, suggesting an involvement of EC in the early genesis of AD^5^. Moreover, we showed that the risk genes enriched for expression in EC overlap substantially with those in microglia (Figure 1D).

Co-expression and two-layer neighbourhood analyses provided insights into some possible functional roles for proteins encoded by AD risk genes expressed in the vascular cells. For example, our results showed lower expression of *PICALM* in EC with AD, suggesting a mechanism by which vascular clearance of Αβ is reduced in AD^27,51^. Functionally less well characterised AD GWAS genes, *WWOX* and *IQCK,* have large neighbourhoods in the FB and PC co-expression networks associated with enrichment for pathways supporting maintenance of BBB integrity^9^. Finally, *INPP5D* had consistently one of the largest two-layer neighbourhoods across the vascular cell types, whereas the enriched pathways associated with them were cell type specific (e.g., EGF/EGFR in EC and ECM components in FB).

Innate immune responses are central to AD pathogenesis and progression but have not been well defined in the microvasculature to date^29,52^. We found evidence for cell-specific differences in vascular inflammatory responses to AD in EC with upregulation of interferon signalling genes in EC. PC also appear to play a role in vascular inflammatory mediation of early AD. Recently identified risk genes *CD46,* encoding a serine protease which mediates inactivation of complement proteins, and *IRAK3,* encoding a homeostatic mediator of innate immune responses^53^ were upregulated and downregulated, respectively, in PC.

However, the most strikingly differentially expressed gene sets in AD are involved in angiogenesis and vascular homeostasis. VEGF/VEGFR and insulin signalling pathways^54^ in EC and EGF/EGFR signalling in EC and PC were downregulated with AD^55^ and negatively correlated with Aβ pathology, despite upregulation of other genes (e.g. *HIF1A)* associated with pro-angiogenic regulation^28^. These results add to prior evidence of *dysfunctional angiogenesis* in AD^16,43,49^. We have extended descriptions by showing that, despite angiogenic signals (e.g., upregulation of *HIF1A*) and metabolic adaptations, major downstream effector pathways fail to respond at the transcriptional level.

Previous reports also have implicated dysfunction of both EC and PC in AD^56^. Our observations emphasise the extent to which pathology of *FB* also contribute to vascular abnormalities in AD, with NOTCH signalling as a candidate mechanism central for this. Multiple genes encoding for ECM components were downregulated in FB, which should contribute to BBB dysfunction^57,58^, impair angiogenesis^35^ and lead to altered expression of tight junction proteins in EC^58^ via reduced signalling between ECM proteins in FB and integrins in EC^59^.

Although we have made a number of novel observations, our analyses had limitations which need to be addressed in future work. Our data was generated from nuclei from multiple brain regions and thus could address robustly only those transcriptomic differences that were common to all of these regions in AD, even given that we took the confound of brain region into account as a fixed effect in the statistical models. A second limitation was that we assessed the total extracted populations of nuclei without seeking to identify and separately study cells expressed from specific vascular anatomic zones^15,17^. Nevertheless, the high overlap of our cellular markers and the cellular markers from human and mouse brain vessel-associated cells in previous reports provides some confidence that all major cell types were represented. Third, like other recent studies, our conclusions are based on relatively sparse (10X Genomics Single Cell 3’ Gene Expression assay) sequencing of the nuclear transcriptome, which may be biased relative to those from the whole cell, potentially reducing the power to detect transcripts from some genes^60^. Use of larger numbers of nuclear and co-expression-based analyses, which rely less on detection of absolute expression levels than do single gene differential expression analyses, may have reduced the impact of this although the impact of this limitation, but this is difficult to assess without future, more comprehensive transcriptional analyses of the whole cells.

Impairment of angiogenesis and vascular homeostasis, reduced endothelial Αβ clearance with reduced expression of *PICALM* and increased production of Αβ by EC with upregulation of interferon (*IFITM3*) and *γ*-secretase component genes all will act to increase toxic Αβ concentrations in the brain^29^. The extraordinary length of the brain capillary network (∼650 km) and its large surface area (∼120 cm^2^/g) suggest that even small relative effects could contribute substantially to increasing the overall Aβ burden in the CNS^61^. The identification of significant EC enrichment in AD risk genes also suggests that their specific contribution to inflammatory activation and reduced Αβ clearance are early, potentially “causal” factors in the onset of sporadic, late onset AD. Our work suggests specific mechanisms by which small vessel disease from many causes (e.g., metabolic disease, hypertension, smoking) could potentiate early AD and add to the rationale for AD prevention through interventions for control of modifiable cardiometabolic risk factors^62^. More generally, our results suggest that EC therapeutic targets related to angiogenic, inflammatory and Αβ clearance pathways deserve prioritisation in the search for treatments able to slow or prevent the onset of AD.

## Methods

Data for this study was generated from cortical brain tissue processed locally as we described earlier^63^ or from that described and made available publicly previously^18^.

### Local tissue access and snRNA sequencing

#### Brain cortical tissue sequencing

Local tissue access and data generation was carried out in accordance with the Regional Ethics Committee and Imperial College Use of Human Tissue guidelines. Tissues and processing were described previously^63^. Cases were selected from the London Neurodegeneration (King’s College London) and Parkinson’s UK (Imperial College London) Brain Banks. Entorhinal and somatosensory cortex from 6 non-diseased control (NDC) cases (Braak stage 0-II) and 6 AD cases (Braak stage III-VI) were used (total of 24 brain samples). Brains used for this study excluded cases with clinical or pathological evidence for small vessel disease, stroke, cerebral amyloid angiopathy, diabetes, Lewy body pathology (TDP-43), or other neurological diseases. Where the information was available, cases were selected for a brain pH greater than 6 and all but one had a *post mortem* delay of less than 24 hr.

### Immunohistochemistry

Immunohistochemical staining was performed on formalin-fixed paraffin-embedded sections from homologous regions of each brain used for snRNASeq locally. Standard immunohistochemical procedures were followed using the ImmPRESS Polymer (Vector Laboratories) and SuperSensitive Polymer-HRP (Biogenex) kits (Table 2). Briefly, endogenous peroxidase activity and non-specific binding was blocked with 0.3% H_2_O_2_ and 10% normal horse serum, respectively. Primary antibodies were incubated overnight at 4°C. Species-specific ImmPRESS or SuperSensitive kits and DAB were used for antibody visualization. Counter-staining for nuclei was performed by incubating tissue sections in haematoxylin (TCS Biosciences) for 2 min. AD pathology was assessed by Aβ plaque (4G8, BioLegend 17-24) and pTau (AT8, NBS Biologics) staining.

**Table 1:**
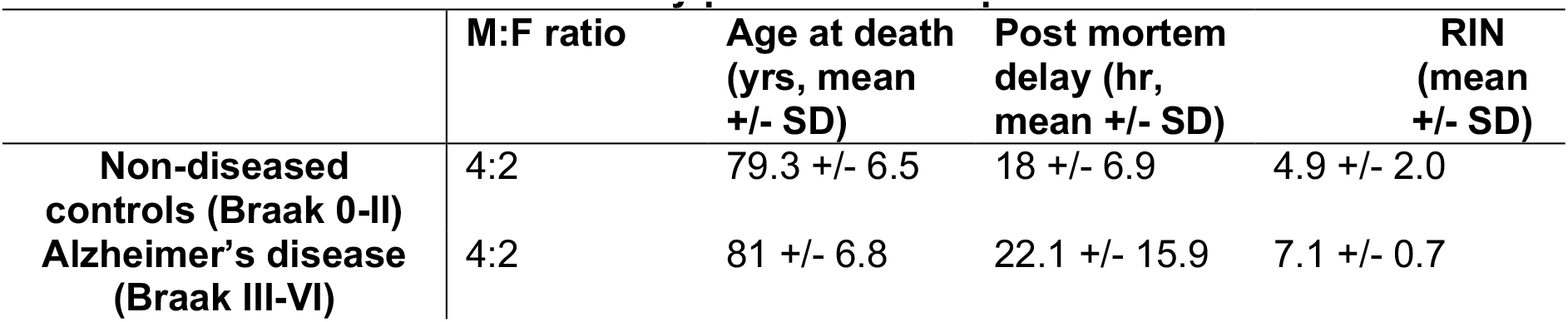
Cohort information for locally processed samples.

**Table 2.**
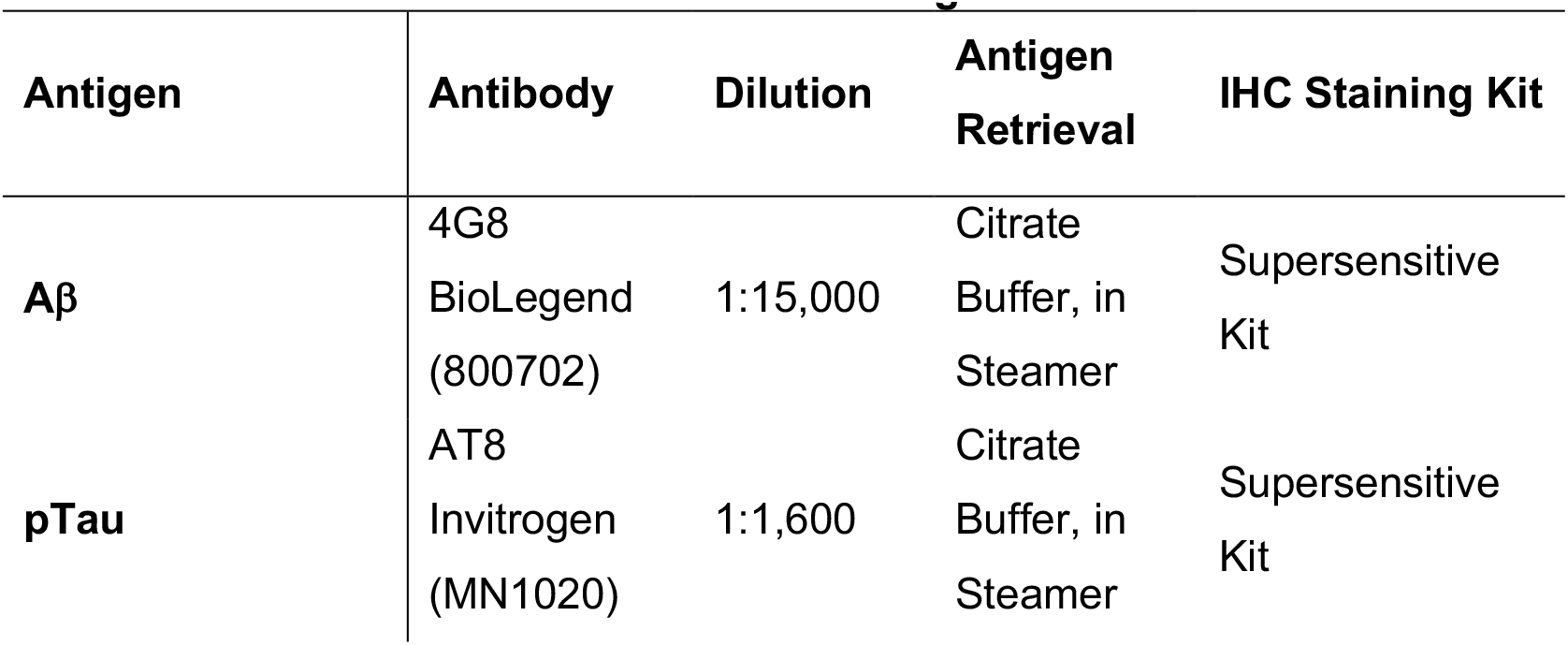
List of antibodies and immunostaining methods.

Labelled tissue sections were imaged using a Leica Aperio AT2 Brightfield Scanner (Leica Biosystems). Images were analysed using HALO software (Indica Labs, Version 2.3.2989.34). The following image analysis macros were used for the study: area quantification macro (amyloid), multiplex macro (pTau).

#### Nuclei isolation and enrichment for lower abundancy cell populations

Local processing of the fresh frozen entorhinal and somatosensory cortical tissue blocks began with sectioning to 80 μm on a cryostat and grey matter separated by scoring the tissue with sharp forceps to collect ∼200 mg grey matter in an RNAse-free Eppendorf tube. Nuclei from NDC and AD samples were isolated in parallel using a protocol based on Krishnaswami *et al.* (2016)^64^. All steps were carried out on ice or at 4°C. Tissue was homogenized in a 2 ml glass douncer containing homogenization buffer (0.1% Triton-X + 0.4 U/μl RNAseIn + 0.2 U/μl SUPERaseIn). The tissue homogenate was centrifuged at 1000 g for 8 min, and the majority of supernatant removed without disturbing the tissue pellet. Homogenised tissue was filtered through a 70 μm filter and centrifuged in an Optiprep (Sigma) density gradient at 13,000 g for 40 min to remove myelin and cellular debris. The nuclei pellet was washed and filtered twice in PBS buffer (PBS + 1% BSA + 0.2 U/ml RNAseIn). Isolated nuclei were labelled in suspension in 1 ml PBS buffer with 1:500 anti-NeuN antibody (Millipore, MAB377, mouse) and 1:250 anti-Sox10 antibody (R&D, AF2864, goat) for 1 hr on ice. Nuclei were washed twice with PBS buffer and centrifuged at 500 g for 5 min. Nuclei were incubated with Alexa-fluor secondary antibodies at 1:1000 (goat-anti-mouse-647 and donkey-anti-goat-488) and Dapi (1:1000) for 30 min on ice, and washed twice. Nuclei were FACS-sorted on a BD Aria II, using BD FACSDiva software, gating first for Dapi +ve nuclei, then singlets and then Sox10- and NeuN-negative nuclei. A minimum of 150,000 double-negative nuclei were collected.

We also isolated nuclei, from adjacent localizations of the same tissue samples as described above, which were not subjected to the FACS-enrichment step but were directly processed for single nucleus capture and snRNA sequencing as described in the following section. In this way, we obtained an unbiased representation of all the brain cell types. The resulting dataset was used for the analysis described in the “Enrichment of brain cell types in AD and WMH GWAS signal” section.

Sorted nuclei were centrifuged at 500 g, resuspended in 50 μl PBS buffer and counted on a LUNA-FL Dual Fluorescence Cell Counter (Logos Biosystems, L20001) using Acridine Orange dye to stain nuclei. Sufficient nuclei were added for a target of 7,000 nuclei for each library prepared. Barcoding, cDNA synthesis and library preparation were performed using 10X Genomics Single Cell 3’ Gene Expression kit v3 with 8 cycles of cDNA amplification, after which up to 25 ng of cDNA was taken through to the fragmentation reaction and a final indexing PCR was carried out to 14 cycles. cDNA concentrations were measured using Qubit dsDNA HS Assay Kit (ThermoFisher, Q32851), and cDNA and library preparations were assessed using the Bioanalyzer High-Sensitivity DNA Kit (Agilent, 5067-4627). Samples were pooled to equimolar concentrations and the pool sequenced across 24 lanes of an Illumina HiSeq 4000 according to the standard 10X Genomics protocol. The snRNAseq data will be made available for download from the Gene Expression Omnibus (GEO) database (https://www.ncbi.nlm.nih.gov/geo/) under accession number GSE160936.

### Single Nuclei RNA Sequence Analyses

#### Processing of FASTQ files

Locally generated snRNASeq data were pre-processed using 10X Genomics Cell Ranger. Illumina sequencing files were aligned to the genomic sequence (introns and exons) using GRCh38 annotation in Cell Ranger v3.1. Nuclei were identified above background by the Cell Ranger software.

#### Quality control, dataset integration, dimension reduction and clustering

Feature-barcode matrices from CellRanger produced corresponding to the local dataset produced as described above were jointly processed with the feature-barcode matrices from a previously published dataset^18^. that were downloaded from the Gene Expression Omnibus (accession number GSE148822). Together, the two datasets were generated from 57 brain samples. Quality control (QC), dataset integration, dimension reduction and clustering were performed using the Nextflow pipeline nf-core/scflow^65^.

QC was performed separately on each sample. Nuclei that had less than 200 features were excluded, whereas for the higher feature filtering criterion, an adaptive threshold was estimated in each sample, which was four median absolute deviations above the median feature number in the sample. Nuclei with more than 5% of mitochondrial gene counts were also excluded. Only genes that had at least one count in 5 nuclei per sample were retained. The QC also included an ambient RNA profiling using the EmptyDrops package^66^ using default parameters. Finally, multiple identification was performed using DoubletFinder^67^ using 10 principal components based on the 2000 most variable features and a pK value of 0.005.

Sample integration was performed using the Liger package (v1.0.0)^19^ incorporated in the nf-core/scflow pipeline (v0.7.1)^65^. The k value was optimized at 20 and the lambda value at 5. 3000 genes were employed in the integration process. The integration threshold was 0.0001 and the maximum number of performed iterations was set to 100. The normalized cell factors from the integrated dataset were then used as an input for dimension reduction and clustering, which was performed using UMAP^20^. Clustering was performed with the Leiden method using a resolution parameter of 0.00001 and a k value of 50.

Cell-type identification of clusters was performed by plotting canonical cell markers using the FeaturePlot function in Seurat (v3.2.3)^68^. To efficiently separate the vascular mural cells, we isolated the ECs and the vascular mural cell clusters and re-ran the steps of the integration, dimension reduction and clustering. Cluster specific genes were identified using the FindMarkers function in Seurat (using the MAST method^25^ with the function arguments set to default). To validate the cell-type specificity of the clusters and their identity, we compared the top 100 cluster markers of our dataset with the top 100 cluster markers of the same cell types from previously published human and mouse datasets^17,21^ using an overrepresentation analysis.

#### Overrepresentation analysis

Overrepresentation analysis was performed to determine if the overlap between two gene sets is significantly higher that if it occurred by chance. This was done using with the “enrichment” function of the R package bc3net (v1.0.4) (https://github.com/cran/bc3net), which performs a Fisher’s exact test (FET). The p values associated with the Fisher’s exact test correspond to the probability that the overlap between the two gene sets and has occurred by chance.

#### Enrichment of brain cell types in AD and WMH GWAS signal

GWAS summary statistics for AD^3^ and WMH (a radiological manifestation of small vessel disease)^24^ were tested for enrichment in brain cell types using the MAGMA.Celltyping (v1.0.1)^5,69^ and MungeSumstats (v1.1.24)^70^ packages. First, summary statistics were appropriately formatted using MungeSumstats for use with MAGMA.Celltyping. Then, SNP associations from the summary statistics were mapped to genes using the map.snps.to.genes function of MAGMA.Celltyping. Next, as described for the default workflow of MAGMA.Celltyping, genes with low variability between the cell clusters were dropped using the drop_uninformative_genes and then quantile groups for each cell type were prepared using the prepare.quantile.groups function.

We first calculated the enrichment in AD and WMH GWAS signal across all the brain cell types on the dataset that had not been subjected to the FACS enrichment step to remove neurons and oligodendrocytes (see “Nuclei isolation and enrichment for lower abundancy cell populations” section) (Figure 1D and S9A, respectively). This was performed using the calculate_celltype_associations function with default parameters and the “linear” enrichment mode. This analysis was repeated after controlling for the microglial enrichment of the GWAS signal. Next, we calculated the enrichment in AD and WMH GWAS signal on each of the vasculature-associated cell types (EC, FB, PC and SMC) (Figure 1E and S9B, respectively). Finally, we assessed if the enrichment of the vasculature-associated cell types in our dataset in AD GWAS signal changed after controlling for the enrichment in WMH GWAS signal. For this, we re-ran the calculate_celltype_associations function for the AD summary statistics and the SNP-to-gene mapping of the WMH GWAS (that was calculated earlier with the map.snps.to.genes function) in the genesOutCOND argument of the function.

#### Differential gene expression analysis

DGE analysis was performed using MAST (v1.18.0). The transcriptomic alterations in AD vs NDC samples were assessed separately in each cell type by means of a zero-inflated regression analysis using a mixed-effects model. The use of a mixed-effects model is particularly important in the context of snRNAseq DGE analyses to account for the pseudoreplication bias that would otherwise be observed if a fixed-effects-only model was employed^71^. The model specification was zlm(∼diagnosis + (1|sample) + cngeneson + pc_mito + sex + brain_region, sca, method = “glmer”, ebayes = F). The fixed effect term pc_mito accounts for the percentage of counts mapping to mitochondrial genes. The term cngeneson is the cellular detection rate. Each nuclei preparation was considered as a distinct sample for the mixed effect. Models were fit with and without the dependent variable and compared using a likelihood ratio test. Units for differential expression are defined as log2 fold difference in AD vs NDC nuclei. The inclusion of a “dataset” term in the model was not necessary because the inclusion of the brain region term completely accounted for it. In the subset of samples that corresponded to the dataset produced in our laboratory, we also performed an exploratory regression analysis of gene expression against the two histopathological features (using pTau or Aβ as markers) using MAST. The model specification was zlm(∼histopath_marker + (1|sample) + cngeneson + pc_mito + sex + brain_region, sca, method = “glmer”, ebayes = F). In this case, units for differential expression are defined as log2 fold difference/% pTau positive cells (or log2 fold difference/% Aβ plaque area), i.e., a one unit change in immunohistochemically-defined pTau (or Aβ plaque) density is associated with one log2-fold change in gene expression. In both MAST analyses, genes expressed in at least 10% of nuclei from each cell type were tested. Genes with an adjusted p-value <0.1 were defined as meaningfully differentially expressed.

#### Gene ontology and pathway enrichment analysis

The gene ontology (GO) enrichment and the pathway enrichments analysis were carried out using the R package enrichR (v 3.0), which uses Fisher’s exact test (Benjamini-Hochberg FDR < 0.1)^72^. Genesets with minimum and maximum genes of 10 and 500 respectively were considered. To improve biological interpretation of functionally related gene ontology and pathway terms and to reduce the number of redundant gene sets, we first calculated a pairwise distance matrix using Cohen’s kappa statistics based on the overlapping genes between the enriched terms and then performed hierarchical clustering of the enriched terms^73^.

#### Gene co-expression analysis

Gene co-expression modules and hub-genes were identified separately for each cell type using the MEGENA (v1.3.7) package^30^. MEGENA constructs a hierarchy of co-expression modules with larger (“parent”) that are further divided into subset (“children”) modules. “Children” modules are subsets of the “parent” ones and have higher numbers as names than their “parents”. To reduce the effect of noise, due to the sparsity of the expression matrix in a snRNAseq experiment, a sample-level pseudo-bulking was performed by summing the raw counts of all the nuclei in a sample. Genes expressed in at least 50% of the samples were used as input. The MEGENA pipeline then was applied using default parameters, using Pearson’s correlations and a minimum module size of 10 genes. Parent modules were produced from which a sub-set of genes form smaller child modules. The co-expression module hierarchy was represented graphically using Cytoscape software (Mac OS version 3.8.0)^74^ (Figure 3A-C).

#### Gene Set Enrichment Analyses (GSEA)

AUCell^75^ (R package v1.6.1) was used to quantify the enrichment of the co-expression modules in our nuclei. Normalised data was processed in AUCell using the *AUCell_buildRankings* function. The resulting rankings, along with the gene lists of interest, were then put into the function *AUCell_calcAUC* (aucMaxRank set to 5% of the number of input genes).

The statistical comparison of the enrichment of the co-expression modules in our AD nuclei vs the NDC nuclei was performed using the limma package in R^31^. The module enrichment matrix was log2-normalized. The default configuration of the limma package was employed with the following linear model (which corresponds to the model employed in the DGE analysis with MAST package): ∼diagnosis+nFeature+pc_mito+brain_region+sex, where nFeatures is the total number of distinct features expressed in each nucleus (to account for the fact that nuclei that express a higher number of features may have higher AUCell scores). We also corrected for a potential pseudoreplication bias^71^, by using the duplicateCorrelation function of the limma package with the sample as the “blocking” variable.

#### Cell-cell communication analysis

Cell-cell communication analysis was performed using CellChat v0.5.5^47^. CellChat employs a curated database of potential signalling ligand-receptor pairs from the literature. Among all these potential ligand-receptor pairs, cell-cell interactions are identified based on mass action models, along with differential expression analysis and statistical tests on cell groups. The CellChat algorithm with default parameters (unless otherwise specified) was applied to the subset of the dataset that corresponded to the ECs, FBs and PCs, separately on the AD and NDC samples. The human CellChat database was used for the ligand-receptor pairs. Communications that involved less that 100 nuclei were filtered out (using the min.cells argument in the filterCommunication function). The ligand-receptor pairs involving genes that showed differential expression in AD compared to the NDC nuclei were identified and are documented in the text as follows: Ligand (expressing celltype(s)) – Receptor (expressing celltype(s)).

## Acknowledgements

We thank the donors and their families for the use of human brain tissue in this study and the London Neurodegeneration and Parkinson’s UK Brain Bank staff for making it available. Infrastructure, including the British Heart Foundation Centre of Excellence, the LMS/NIHR Imperial Biomedical Research Centre Flow Cytometry Facility and the Imperial BRC Genomics Facility, which are supported by the National Institute for Health Research (NIHR) Biomedical Research Centre (BRC), all contributed to the work. ST was supported by an “Early Postdoc.Mobility” scholarship (P2GEP3_191446) from the Swiss National Science Foundation, a “Clinical Medicine Plus” scholarship from the Prof Dr Max Cloëtta Foundation (Zurich, Switzerland) and a scholarship from the University Hospitals of Geneva. PMM acknowledges generous personal support from the Edmond J Safra Foundation and Lily Safra and an NIHR Senior Investigator Award. This work was supported by the UK Dementia Research Institute, which receives its funding from UK DRI Ltd., funded by the UK Medical Research Council, Alzheimer’s Society and Alzheimer’s Research UK.

## Data availability

The snRNAseq data will be made available for download from the Gene Expression Omnibus (GEO) database (https://www.ncbi.nlm.nih.gov/geo/) under accession number GSE160936. Previously described data^18^ was downloaded from the GEO database (GSE148822).

## Author Contributions

Stergios Tsartsalis – experimental design, primary data analysis, interpretation of results, drafted the manuscript

Nurun Fancy – experimental design, primary data analysis, interpretation of results

Amy Smith-performed experiments, data analysis, interpretation of results

Karen Davey – performed experiments, interpretation of results

Combiz Khozoie – software developments, support for data analysis

Xin Ye – primary regression analyses, interpretation of results

Nanet Willumsen – performed immunohistochemistry experiments and image analysis

Aisling McGarry-performed immunohistochemistry experiments and image analysis

Robert C.J. Muirhead-performed immunohistochemistry experiments and image analysis

Stephanie Debette – supported data analysis, interpretation of results

David Owen – experimental design, interpretation of results

Paul M. Matthews-experimental design, interpretation of results, drafted the manuscript

All of the co-authors reviewed and approved the final manuscript.

## Competing Interests

PMM has received consultancy fees from Roche, Adelphi Communications, Celgene, Neurodiem and Medscape. He has received honoraria or speakers’ fees from Novartis and Biogen and has received research or educational funds from Biogen, Novartis and GlaxoSmithKline.

---

**Figure S1.**
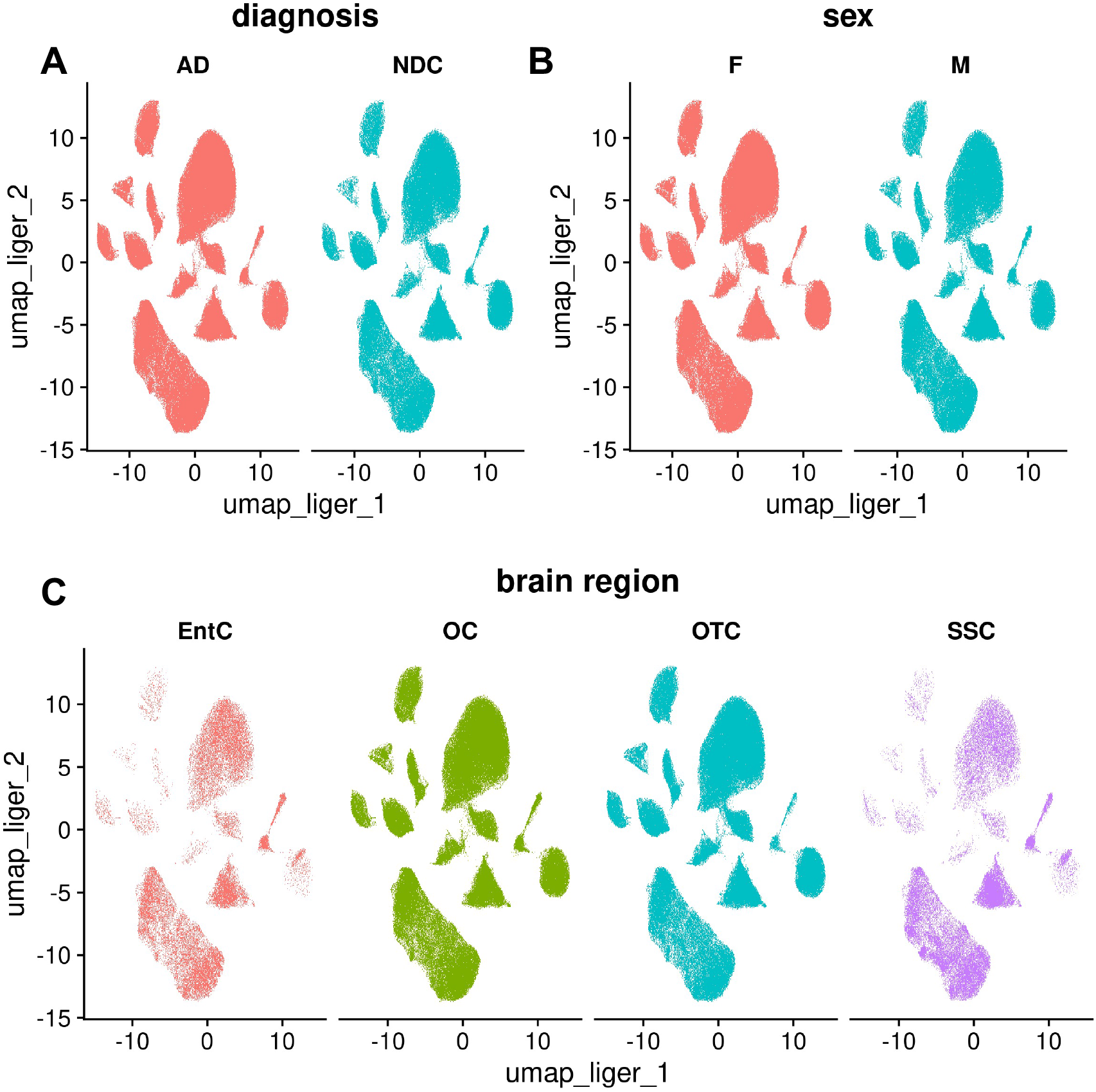
UMAP plots of the integrated snRNAseq dataset (see Figure 1A) by diagnosis (A), sex (B) and brain region. (EntC, entorhinal cortex; OC, occipital cortex; OTC, occipital temporal cortex; SSC, somatosensory cortex) showing that the nuclei were well mixed with respect to these parameters after integration

**Figure S2.**
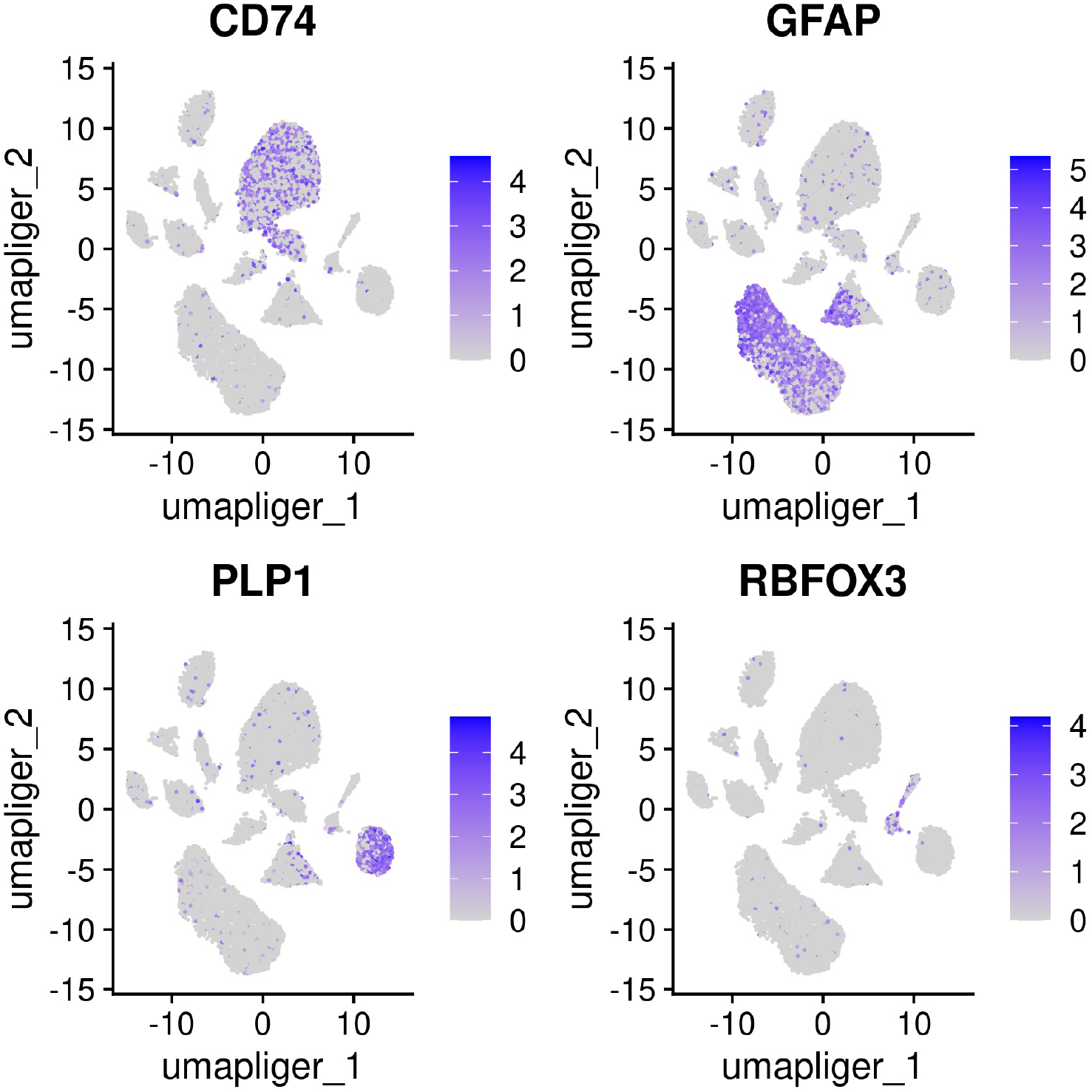
UMAP feature plots of canonical cell marker genes for microglia (*CD74*), astrocytes (*GFAP),* oligodendrocytes (*PLP1)* and neurons (*RBFOX3)*.

**Figure S3.**
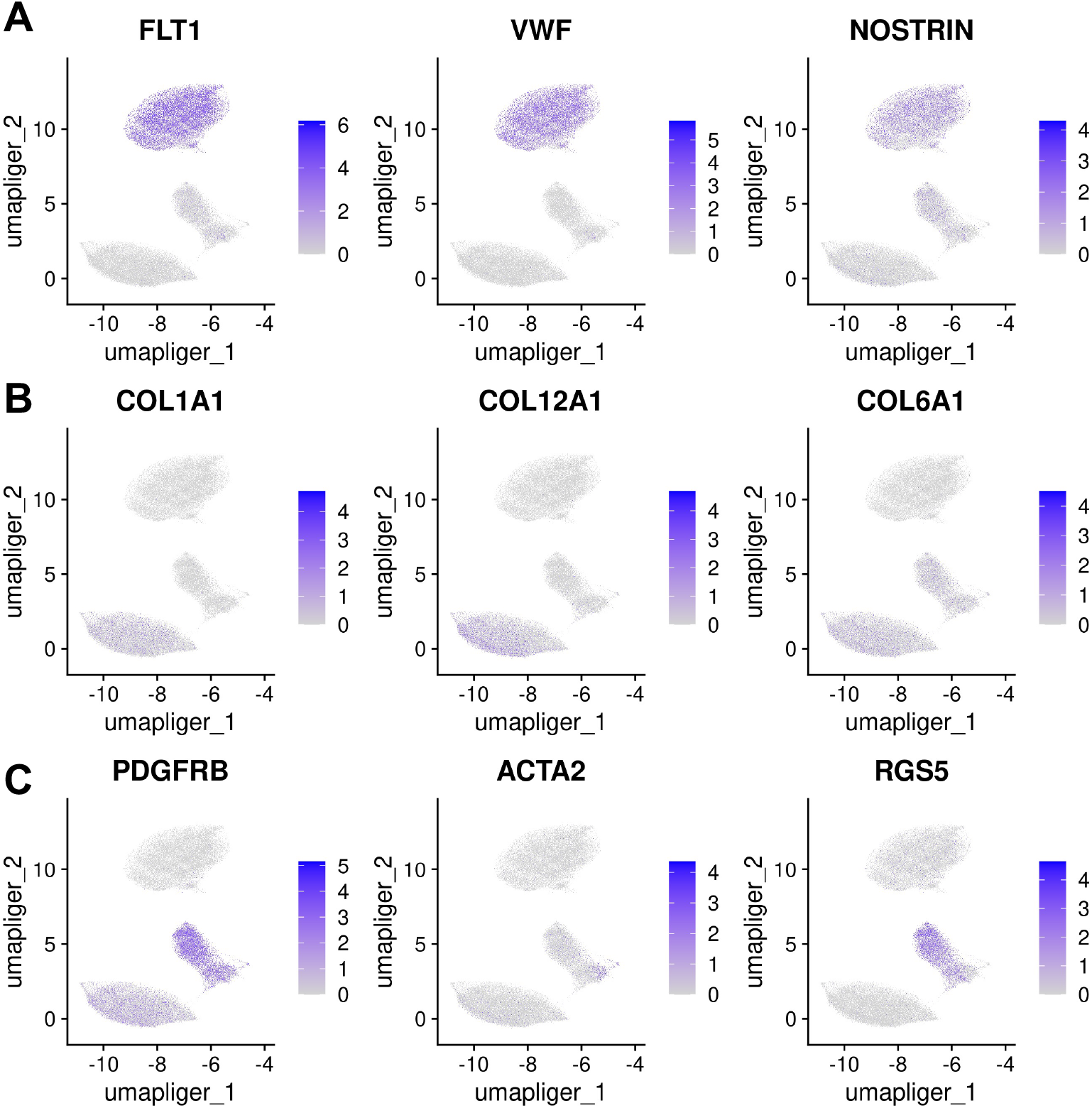
UMAP feature plots of marker genes for EC (A), FB (B) and PC and SMC (C).

**Figure S4.**
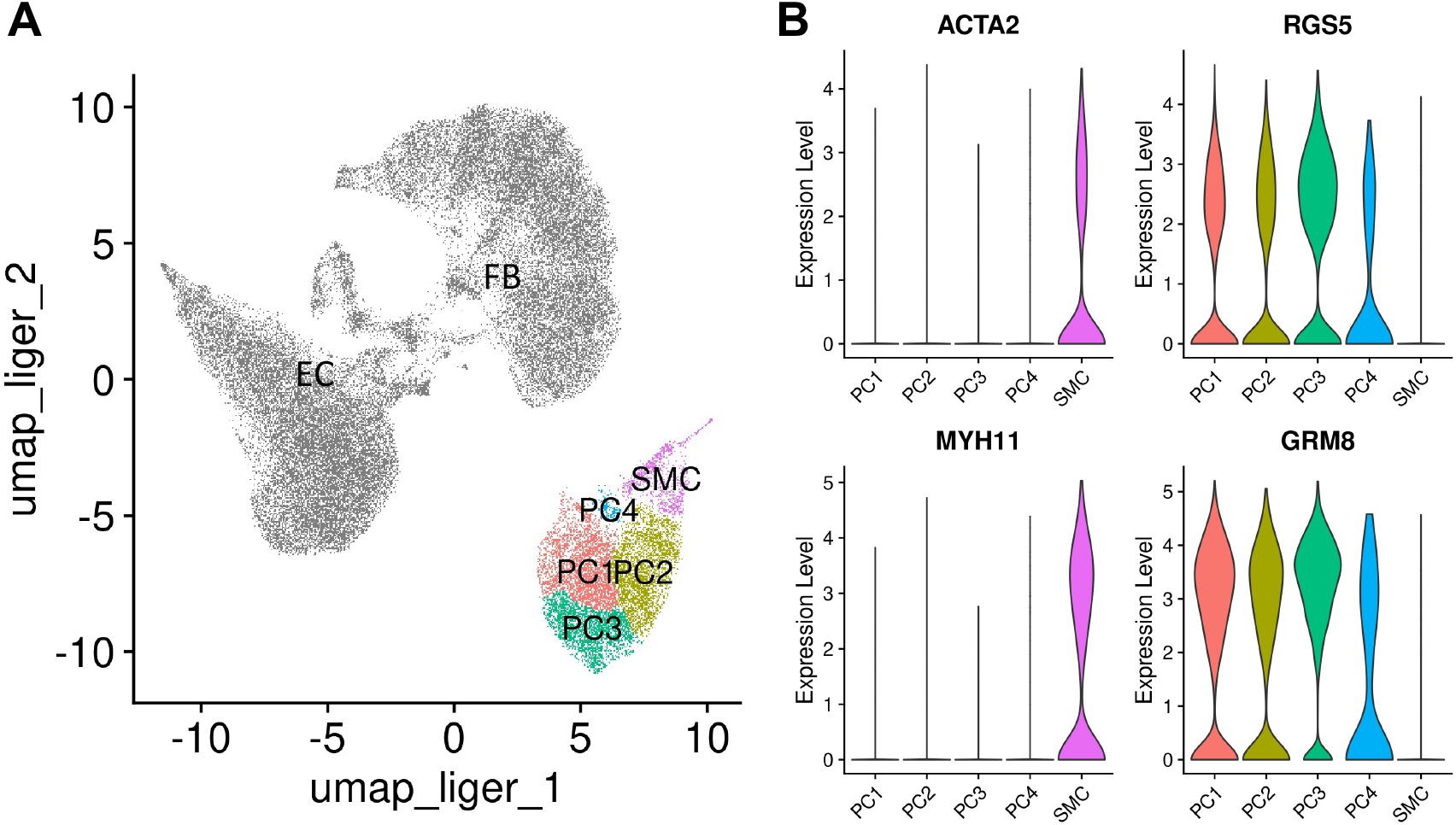
UMAP plot after re-integration and clustering of the EC, FB, PC and SMC nuclei in the integrated dataset. (A). EC and FB nuclei are coloured in grey. Four subclusters (PC1, PC2, PC3, PC4 and SMC) correspond to PC and SMC (B). Violin plots of genes previously shown to be specific for PC (*RGS5* and *GRM8)* and SMC *(ACTA2, MYH11)*.

**Figure S5.**
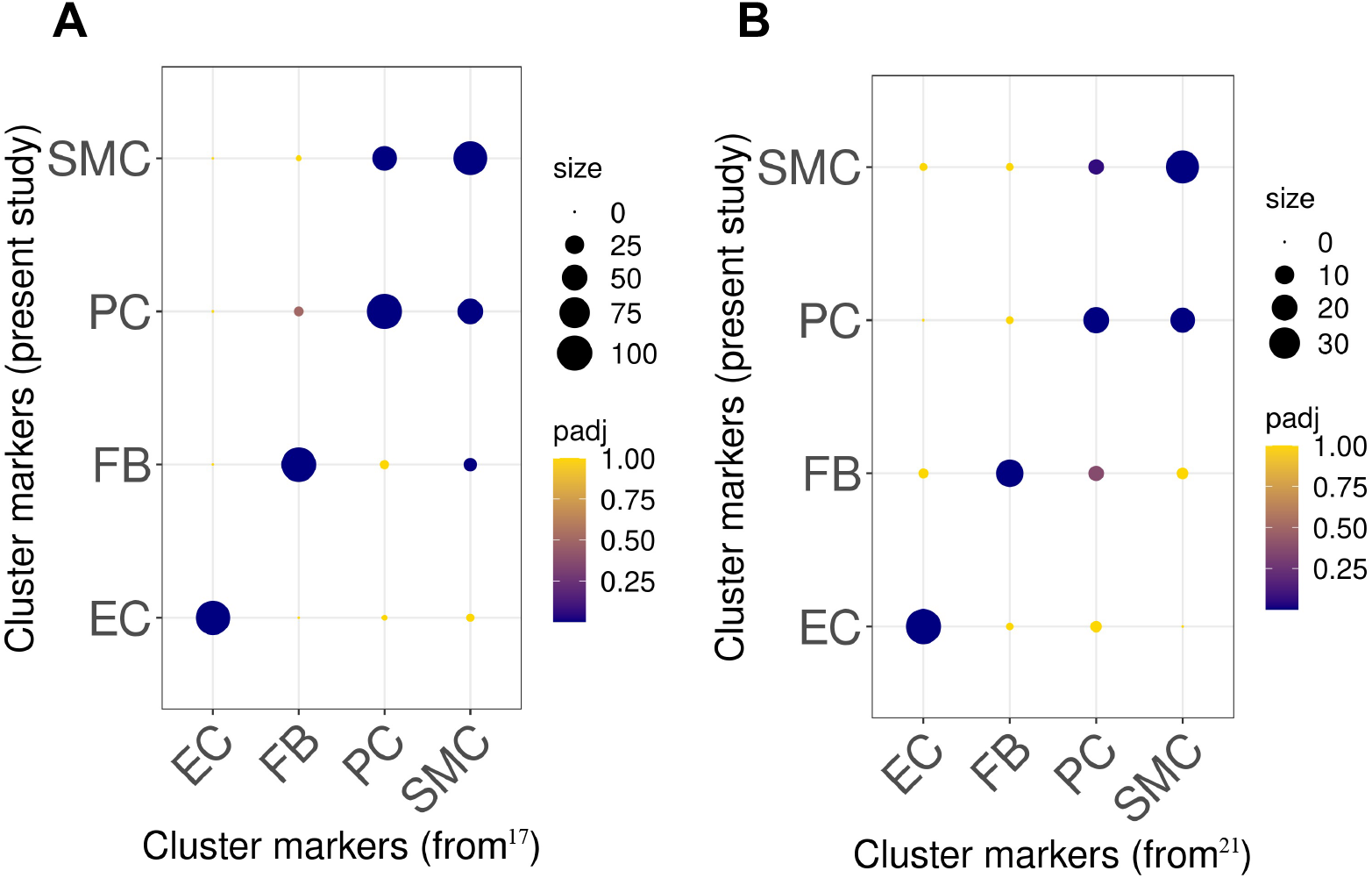
Dot plots of the overlap between cell markers for EC, FB, PC and SMC previously identified for (A) human^17^ and (B) mouse^21^ and the cluster markers used in the present study. The size of the dots correspond to the overlap between the cluster gene sets and the colour of the dot to the adjusted p-value of an overrepresentation Fisher’s exact test.

**Figure S6.**
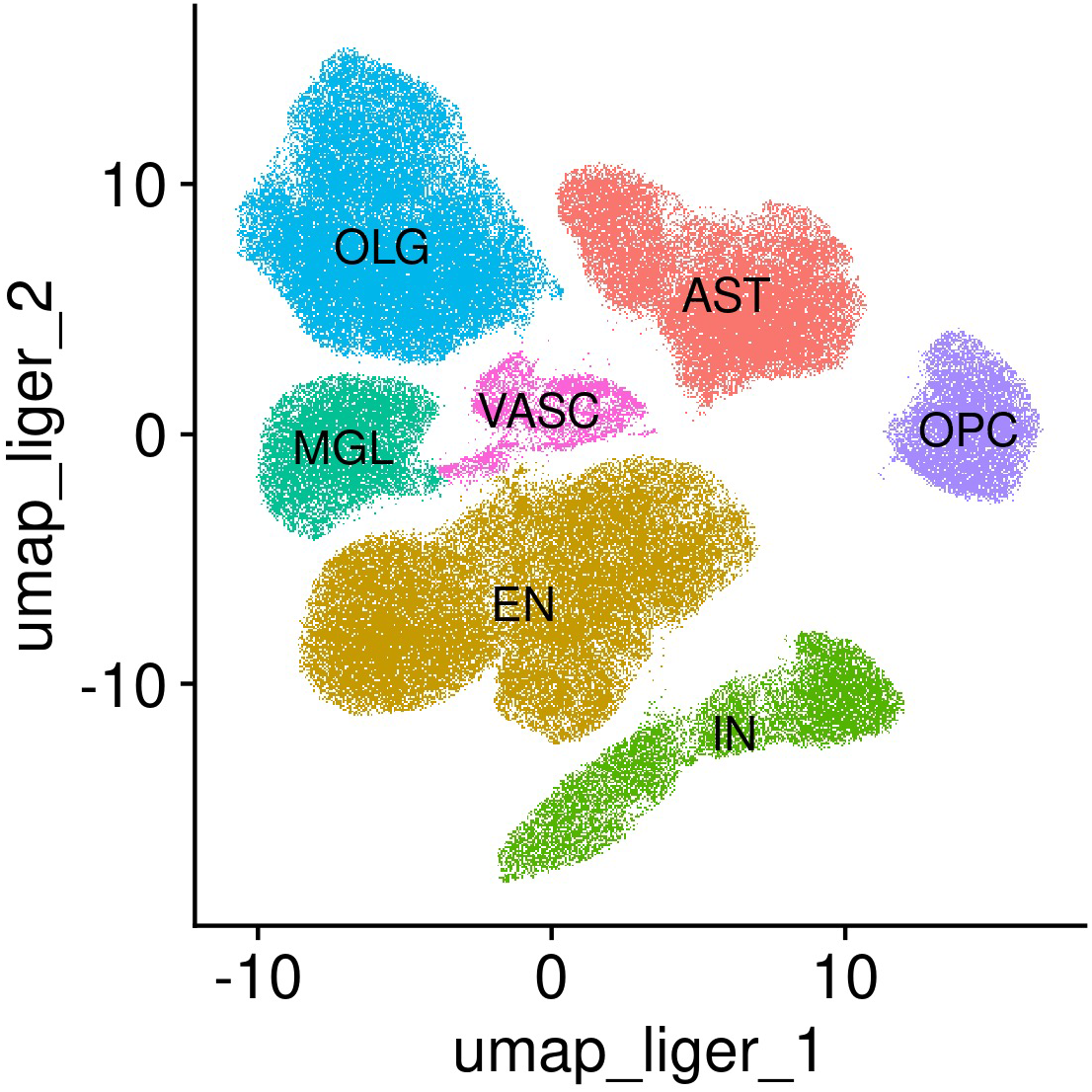
UMAP plot of the snRNAseq dataset that was generated without prior FACS-enrichment step to remove neurons and oligodendrocytes. AST, astrocytes; EN, excitatory neurons; IN, inhibitory neurons; MGL, microglia; OLG, oligodendrocytes; OPC, oligodendrocyte progenitor cells; VASC, vascular cells.

**Figure S7.**
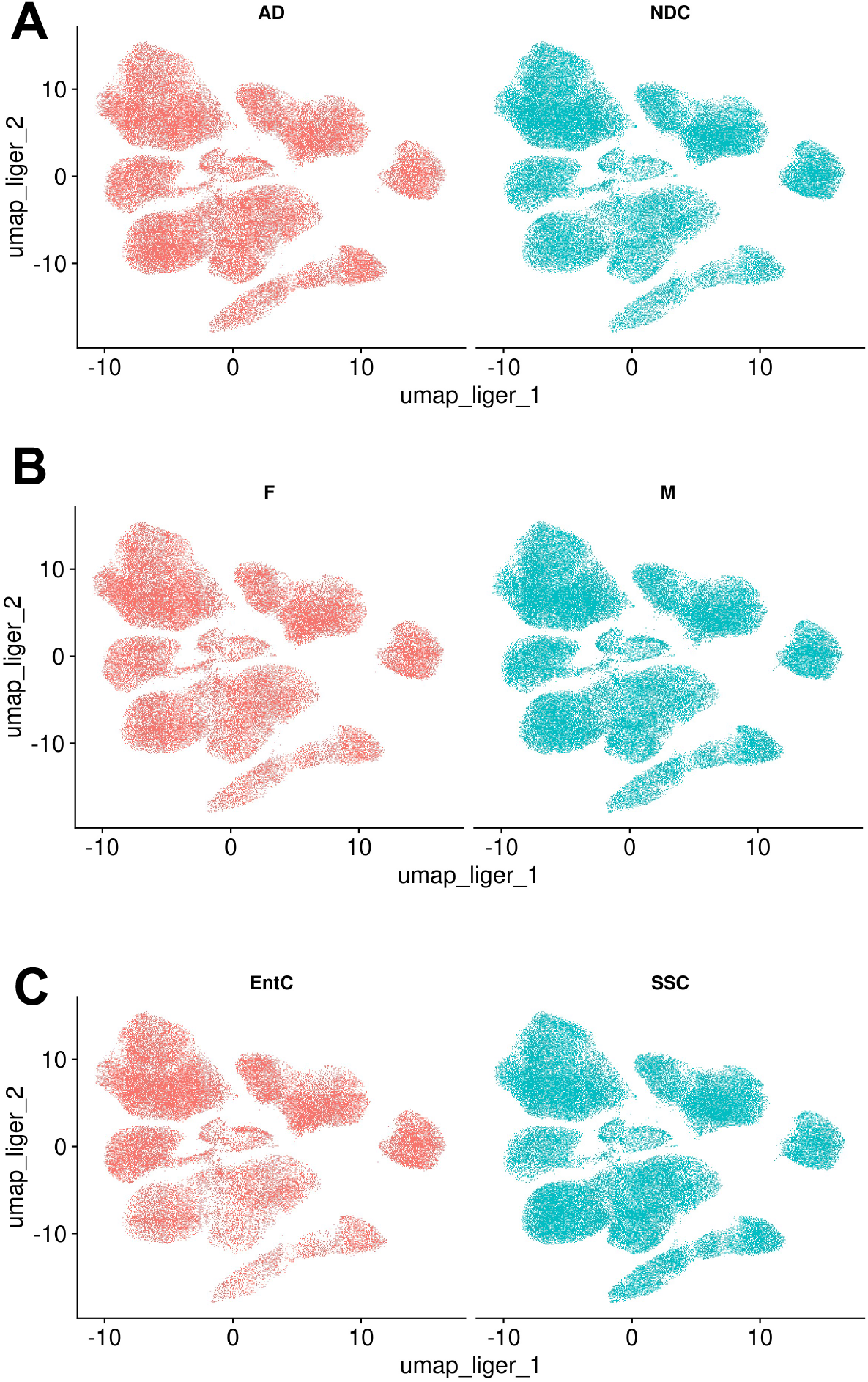
UMAP plots of the snRNAseq dataset that was generated without prior FACS-enrichment step (Figure S6) by diagnosis (A), sex (B) and brain region. (EntC, entorhinal cortex; OC, occipital cortex; OTC, occipital temporal cortex; SSC, somatosensory cortex), showing that the nuclei were well mixed with respect to these parameters after integration.

**Figure S8.**
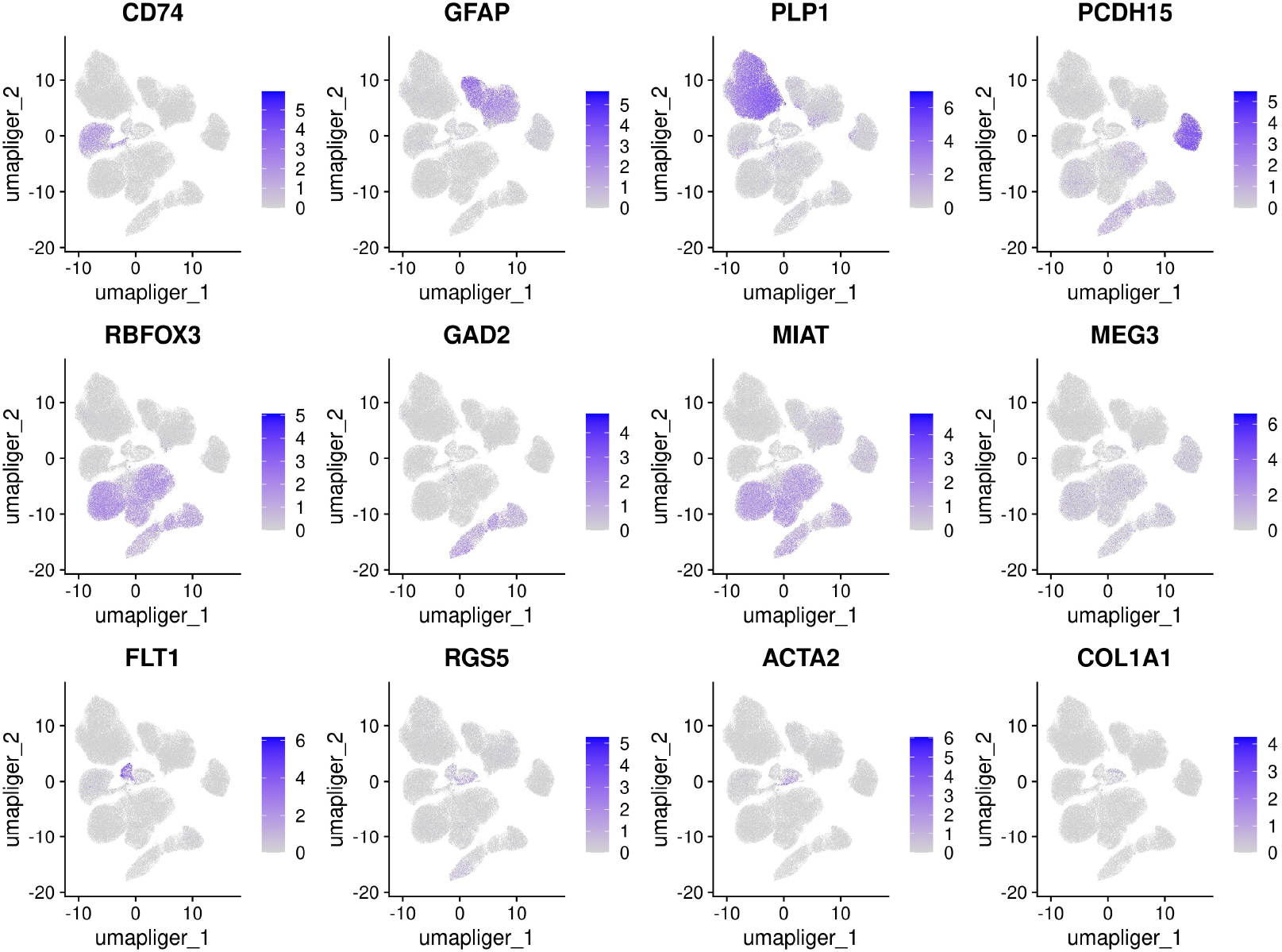
UMAP feature plots of canonical cell marker genes in the dataset of Figure S6 and S7 for microglia (*CD74*), astrocytes (*GFAP),* oligodendrocytes (*PLP1),* oligodendrocyte precursor cells (*PCDH15*), neurons (*RBFOX3, GAD2, MIAT, MEG3*) and vascular cells (*FLT1, RGS5, ACTA2, COL1A1)*.

**Figure S9.**
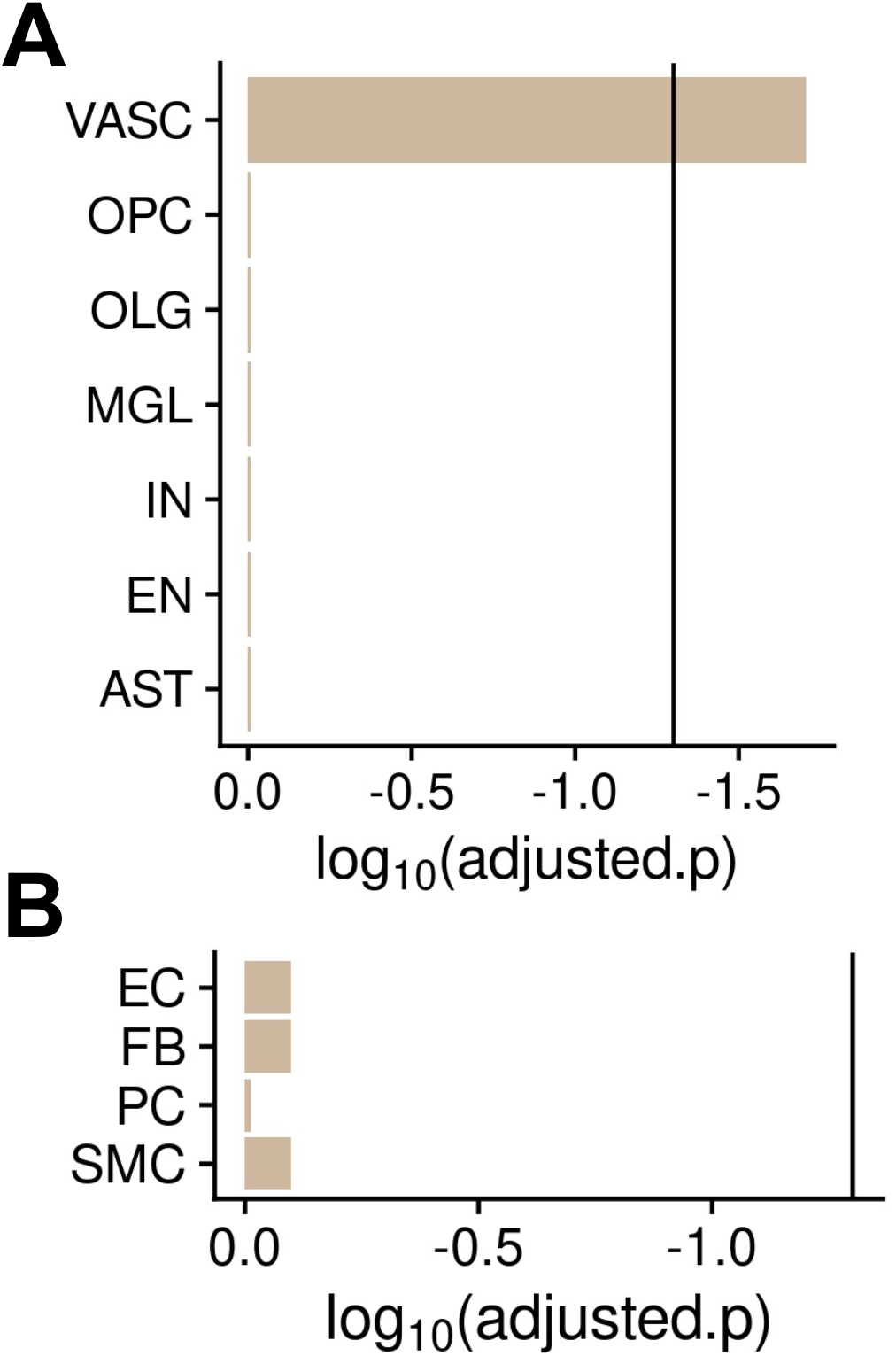
MAGMA.Celltyping enrichment of brain nuclei in genomic loci associated with genetic risk for WMH. The bars correspond to the log_10_ p value of the enrichment. The dark line marks the corrected significance threshold. Only vascular nuclei show enrichment with fully corrected significance across the whole dataset. (F) MAGMA.Celltyping WMH risk gene enrichment of nuclei of the brain vasculature showing no significant enrichment for any of the cell types, suggesting that the enrichment for genomic loci associated to WMH is equally distributed among them.

